# 600+ insect genomes reveal repetitive element dynamics and highlight biodiversity-scale repeat annotation challenges

**DOI:** 10.1101/2022.06.02.494618

**Authors:** John S. Sproul, Scott Hotaling, Jacqueline Heckenhauer, Ashlyn Powell, Dez Marshall, Amanda M. Larracuente, Joanna L. Kelley, Steffen U. Pauls, Paul B. Frandsen

## Abstract

Repetitive elements (REs) are integral to the composition, structure, and function of eukaryotic genomes, yet remain understudied in most taxonomic groups. We investigated REs across 601 insect species and report wide variation in REs dynamics across groups. Analysis of associations between REs and protein-coding genes revealed dynamic evolution at the interface between REs and coding regions across insects, including notably elevated RE-gene associations in lineages with abundant long interspersed nuclear elements (LINEs). We leveraged this large, empirical data set to quantify impacts of long-read technology on RE detection and investigate fundamental challenges to RE annotation in diverse groups. In long-read assemblies we detected ∼36% more REs than short-read assemblies, with long terminal repeats (LTRs) showing 162% increased detection, while DNA transposons and LINEs showed less respective technology-related bias. In most insect lineages, 25–85% of repetitive sequences were “unclassified” following automated annotation, compared to only ∼13% in *Drosophila* species. Although the diversity of available insect genomes has rapidly expanded, we show the rate of community contributions to RE databases has not kept pace, preventing efficient annotation and high-resolution study of REs in most groups. We highlight the tremendous opportunity and need for the biodiversity genomics field to embrace REs and suggest collective steps for making progress towards this goal.

## Introduction

Repetitive elements (REs) comprise large proportions of eukaryotic genomes and are fundamental to the evolutionary process (Bourque et al. 2018; Gilbert et al. 2021). Broadly, REs can be classified as *interspersed* or *tandem repeats*. Interspersed repeats include transposable elements (e.g., retrotransposons) which encode for proteins that facilitate their movement and proliferation in genomes. Tandem repeats (e.g., satellite DNAs) can form large blocks (e.g., megabases) of relatively short non-coding sequences in repeated arrays (reviewed in UgarkoviĆ and Plohl 2002). Together, interspersed and tandem repeats comprise ∼67% of the human genome (de Koning et al. 2011). Despite their major genomic footprint, REs are understudied in genome science due to a history of technical challenges associated with their sequencing and assembly (Bergman and Quesneville 2007; Sotero-Caio et al. 2017); however, long-read sequencing is ameliorating this challenge through improvements in genome assembly contiguity (Hotaling et al. 2021b).

Although understudied, REs can play critical roles in the organization, stability, regulation, and evolution of genomes (Bourque et al. 2018; Wells and Feschotte 2020). At broad scales, REs shape chromatin domains across chromosomes and impact the three-dimensional organization of DNA (Sun et al. 2020; Winter et al. 2018). Rapidly evolving blocks of REs are common sites of recombination and chromosome rearrangements (e.g., Bennetzen 2000; Cáceres et al. 1999). At finer scales, shifts in RE location and abundance (e.g., through transposition of retrotransposons) can alter gene expression and phenotype evolution (Chuong et al. 2017; Schrader and Schmitz 2019; Stuart et al. 2016). Across evolutionary scales, rapid RE evolution (e.g., tandem repeats) is associated with hybrid incompatibilities between species (Brand and Levine 2022; Ferree and Barbash 2009; Jagannathan and Yamashita 2021). In short, REs exhibit an array of structural and evolutionary effects on genome evolution across species.

We have entered the era of biodiversity genomics with availability and quality of genome assemblies rapidly increasing in plants and animals (Hotaling et al. 2021b, 2021a; Marks et al. 2021). As a critical mass of assemblies accumulates within a group, phylogenetically informed meta-analyses of REs can illuminate their impact on genome dynamics and evolution (Gilbert et al. 2021). With more than 1 million described species, insects account for the bulk of Earth’s animal biodiversity (Stork 2018). While some insects have become model genetic organisms (e.g., *Drosophila melanogaster*) and thus considerable attention is devoted to many aspects of their genome biology, including REs (e.g., Kim et al. 2014; Vargas-Chavez et al. 2022), for the vast majority of insects (and many other taxonomic groups), repetitive genomic components remain largely unexplored.

In this study, we analyzed the RE landscape in genome assemblies of more than 600 insect species that have diverged from common ancestors over ∼400 million years of evolution (Misof et al. 2014). We used this dataset to gain a broad view of RE dynamics in insects and to assess how sequencing technology and taxonomic representation in reference databases (e.g., Repbase, Bao et al. 2015) shapes our ability to identify and classify REs with widely used automated RE annotation tools in a comparative framework. Given the potential for REs to impact protein-coding genes (e.g., through epigenetic silencing of adjacent DNA sequences), we also investigated the frequency of associations between REs and protein-coding genes. An inherent challenge to broad-scale analysis of publicly available assemblies is that variation in the finer technical details (e.g., specific sequencing platform used, assembly method) can lead to variation in per-assembly quality and add noise to results. However, these caveats can be balanced by the potential for detecting broad-scale trends in signal that are only obvious when sampling many species within and across lineages. We reduce the impacts of per-assembly quality by filtering the lowest quality assemblies and identify robust, broad-scale trends by visualizing RE dynamics across hundreds of species in their phylogenetic context. Our findings yield new insight into the RE landscape of insect genomes from a much wider taxonomic perspective than previous analyses and identify trends that spawn new hypotheses surrounding the role of REs on shaping genome evolution within lineages. Beyond insects, we use this large, diverse data set to highlight the opportunities and obstacles for investigating RE dynamics in biodiversity studies with emphasis on RE annotation bottlenecks. We conclude by describing the ways in which the biodiversity genomics community can alleviate challenges of RE annotation (e.g., RE database curation and taxonomy) to build towards a more holistic understanding of genomic natural history and evolution.

## Results

We assessed RE content for genome assemblies of 601 insect species across a total of 20 orders. Of the 601 assemblies, 548 and 441 assemblies had BUSCO completeness ≥ 50% and ≥ 90%, respectively. We report results for three data sets: all assemblies, ≥ 50% BUSCO, and ≥ 90% BUSCO. For clarity, we used the “all assemblies” data set for analyzing taxonomic representation in Repbase, the “≥ 50%” data set for assessing overall RE trends in insects, and the “≥ 90%” for all other analyses.

The proportion of REs in insect genomes ranged widely from 1.6–81.5% (mean = 30.8%; Figs. 1 & 2). Based on mean genomic proportion of specific RE categories, DNA transposons were the most abundant overall and particularly so in Coleoptera (Figs. 1, 2a–d), yet conspicuously uncommon in Lepidoptera. LINEs were the next most abundant RE type and exhibited wide variance across and within orders (Figs. 1, 2a–d). For example, within Diptera, LINE abundances ranged from ∼1% in 29 species to ∼47% in *Hermetia illucens* (Fig. 1b). However, LINEs were notably uncommon in Hymenoptera (1.8% ± 1.7% genomic proportion, n = 157 species). LTRs were generally uncommon but were abundant within *Drosophila* (order Diptera; Fig. 1b). Since LTRs are particularly difficult to identify due to their size and complexity (Flynn et al. 2020), and because *Drosophila* LTRs are better characterized in RE databases than other insect lineages (see below), this trend may reflect methodology more than biology. Consistent with previous studies (Cong et al. 2022; Heckenhauer et al. 2022; Petersen et al. 2019), we showed that RE abundance correlates with genome size (Fig. 2j). Previous studies (Schley et al 2022; Novak et al 2020) have noted an inflection point in this correlation where increasingly large-genome (e.g.,>5–10 Gb) species have lower than predicted genomic proportion of repeats, likely because remains of retained ancient TE bursts accumulate sufficient mutations that RE detection software classifies them as unique/low-copy sequences rather than repetitive sequences. We did not find evidence of such an inflection point in insects (Supplemental Fig. S3a), however we note that our data set includes a poor sampling of large-genome species (e.g., few assemblies exceed 1 Gb and none exceed 5 Gb). SINEs showed greatest abundance in Blattodea, Phasmatodea, Lepidoptera and some dipterans, tandem repeats were most common in Hymenoptera and Diptera, while “Other” repeats were especially abundant in Lepidoptera reflecting the high number of Helitrons in some lineages (Supplemental Figs. S1 and S3b–d).

**Figure 1.**
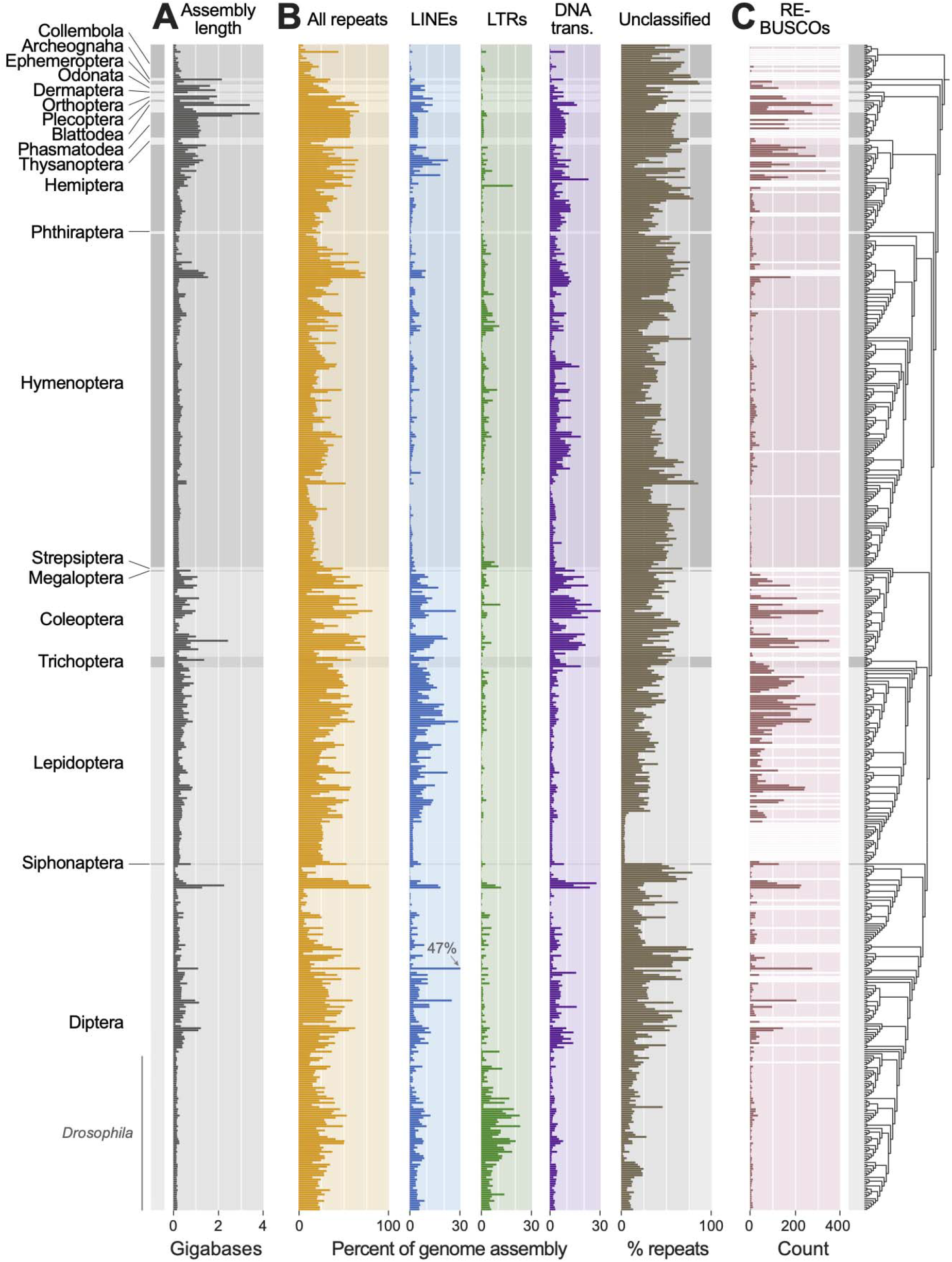
The repetitive element landscape of insects. Left bars with alternating shades of gray indicate taxonomic boundaries and track across plots. (A) Total genome assembly length. (B) Overall RE abundance followed by LINEs, LTRs, and DNA transposons all as a percent of the overall genome assembly. Totals for DNA transposons reported include TIR, Crypton, and Helitron/Polinton elements, reflecting the classification scheme of RepeatMasker, although the large majority in all cases are TIR – see Supplemental Fig. S2 for a finer breakdown of these three categories. One species (*Hermetia illucens*) exceeded the scale for LINEs (indicated at 47%). Any REs that could not be classified (“unclassified”) are shown as a percentage of all repeats identified for a given species. (C) Abundance of RE-associated BUSCOs. For (A) and (B), all assemblies with BUSCO completeness ≥ 50% were included (n = 548). For (C), because we were concerned that BUSCO completeness would alter our capacity to detect RE-associated BUSCOs, we only included assemblies with BUSCO completeness ≥ 90% (n = 493). Summary of less abundant repeat categories is shown in Supplemental Fig. S1, Supplemental Materials. Assemblies that were excluded in (C) are indicated with white bars. To the right of the plots, the phylogeny inferred in this study that was used to place species in their phylogenetic context is shown.

**Figure 2.**
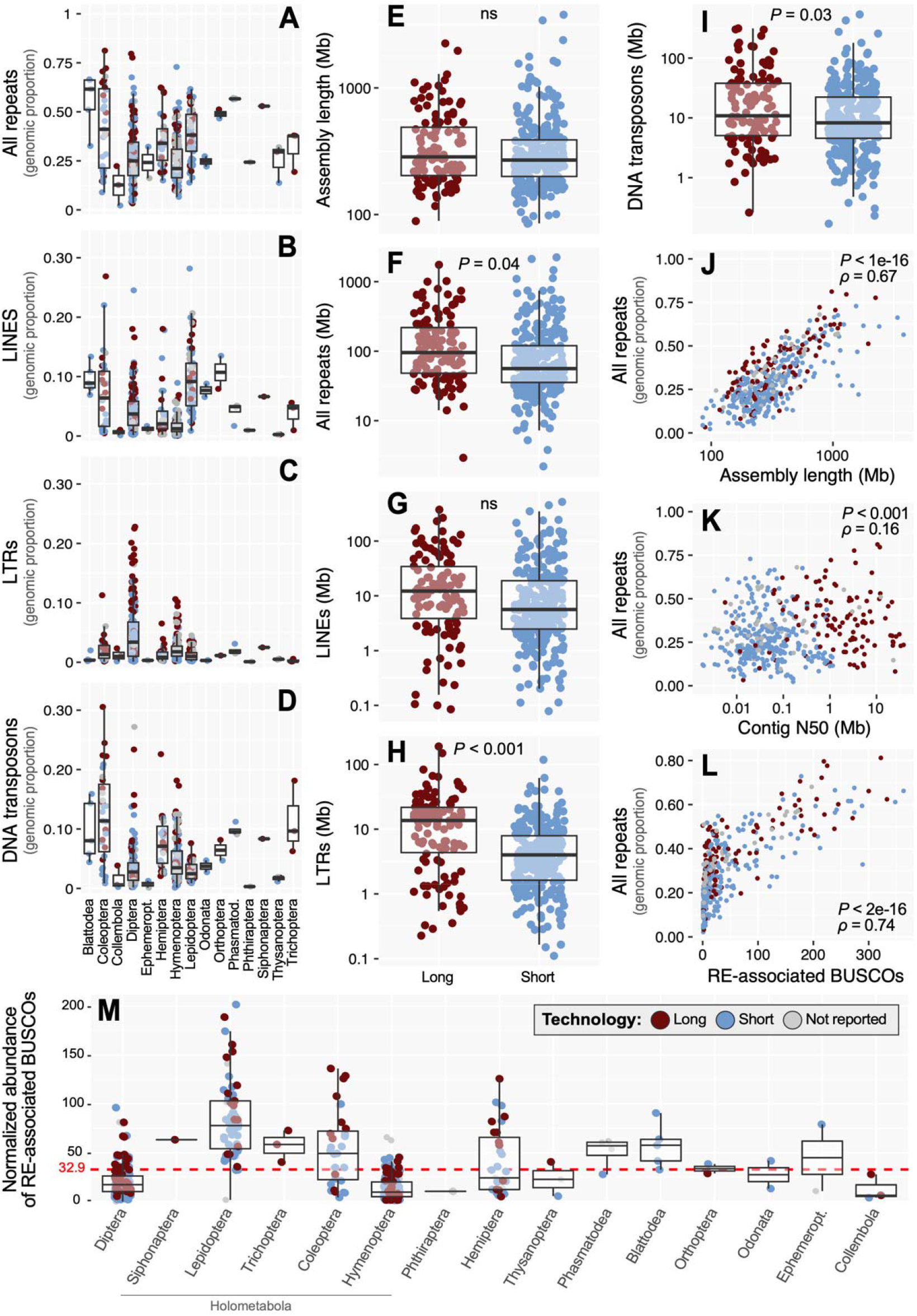
Statistical summaries of insect repetitive element dynamics, technology impacts. (A–D) Genomic proportion of all repeats (A), LINEs (B), LTRs (C), and DNA transposons (D) across insect orders in the data set. Note: To improve visualization, y-axis scales differ between (A) and (C–D). (E–I) Sequencing technology comparisons for (E) assembly length, (F) all repeats, (G) LINEs, (H) LTRs, and (I) DNA transposons. Totals for DNA transposons reported include TIR, Crypton, and Helitron/Polinton elements, reflecting the classification scheme of RepeatMasker, although the large majority in all cases are TIR – see Supplemental Fig. S2 for a finer breakdown of these three categories. Significance was assessed with Welch two-sample *t*-tests. ns: not significant at *P* < 0.05. (J–L) Spearman correlations between genomic proportion of repeats versus (J) assembly length, (K) contig N50, (L) and number of RE-associated BUSCO genes. (M) Normalized abundances of RE-associated BUSCOs across orders and organized by the phylogeny shown in Fig. 1. For all plots, log-transformed data were used for visualization while statistics were performed on the untransformed data.

Comparison of assembly-based (AB) estimates of RE abundance to assembly-free clustering-based (CB) estimates in dnaPipeTE (Goubert et al. 2015) showed that patterns of relative abundance in RE categories were broadly consistent across methods (Supplemental Fig. S4). Our second CB approach using RepeatExplorer2 (RE2) (Novák et al. 2013) also showed general consistency in patterns, albeit with low resolution in the classification of interspersed repeats – a pattern that has been noted in other insect studies and is likely related to the general Metazoa library used by RE2 being poorly suited to insect annotation (Heckenhauer et al. 2022). Both CB approaches consistently showed higher estimates of tandem repeats than the AB estimates (Supplemental Figs. S4 and S5), which corroborates the expectation that blocks of highly similar repeats are underrepresented in assemblies (Novák et al. 2010). When comparing overall repeat proportion estimates across methods, two patterns emerged. In seven of 15 comparisons, overall repeat proportion estimates showed minor variation such that genomic proportion of repeats varied only 1–10% across methods (e.g., see Nicrophorous, Aethinia, and Aphidius, Supplemental Fig. S4). In the remaining comparisons (i.e., eight of 15), the genomic proportion of repeats in the AB approach differed by >15% when compared to one or both CB estimates. In all but one of these cases, the AB approach estimated a notably higher proportion of repeats than either of the CB approaches, which is counter to the notion that assembly-based approaches often underestimate repetitiveness. A correlated characteristic in this subset of samples is that in each case, the AB analysis found notably high proportions of interspersed repeats (e.g., see DNA Transposons + LINEs in Limonius, Harmonia, and Bemisia, Supplemental Fig. S4). Taken together, our findings both corroborate the improved estimates of certain repeat classes such as tandem repeats expected by CB approaches, and present evidence that the same CB approaches can be prone to large underestimates of interspersed repeats compared to long-read AB approaches.

Sequencing technology influences the study of REs. In insects, long-read assemblies are on average ∼48 times more contiguous than short-read technologies (Hotaling et al. 2021b). For REs, we identified 36.1% more REs in long-read assemblies versus short-reads (Welch two-sample *t*-test, *P* = 0.04; Fig. 2f). Furthermore, this difference in total REs identified was not due to assembly length, which did not differ between technologies (Welch two-sample *t*-test, *P* = 0.42; Fig. 2e), however a positive correlation between repeat abundance and assembly contiguity (i.e., contig N50) was observed (Welch two-sample *t*-test, *P* = >0.001; Fig. 2k). Long-read assemblies had the greatest influence on LTR detection (162% increase, Welch two-sample *t*-test, *P* < 0.001; Fig. 2h), followed by DNA transposons (47% increase; Welch two-sample *t*-test, *P* = 0.03; Fig. 2i). Although LINEs showed increased average detection in long-read assemblies the difference was not significant (Welch two-sample *t*-test, *P* = 0.42; Fig. 2g).

These trends set a general expectation for sequencing technology-related bias, with LTRs being under-detected in short-read assemblies, whereas DNA transposons, LINEs, and other RE classes, show moderate/low sensitivity to sequencing technology in assembly-based RE detection (Fig. 2f-i). As a surrogate measure for associations between REs and protein-coding genes, we quantified RE presence in BUSCO genes (termed hereafter RE-associated BUSCOs) following Heckenhauer et al. (Heckenhauer et al. 2022). RE-associated BUSCOs increased with overall repeat content in assemblies (Spearman’s correlation = 0.74, *P* < 2 x 10^16^; Fig. 2l). However, assembly repeat content alone did not explain increased RE-associated BUSCO abundance. For example, Lepidoptera and Coleoptera species had 5.8- and 4.4-fold respective average increases in RE-associated BUSCOs compared to Hymenoptera after correcting for assembly length (Fig. 2m). Overall, RE-associated BUSCOs were most abundant in species with high proportions of LINEs (e.g., Hemiptera, Blattodea, Coleoptera, Trichoptera, Lepidoptera) (Figs. 1, 2m, and Supplemental Fig. S6). In some lineages (e.g., some Blattodea, Coleoptera, and Hemiptera), RE sequences were detected in upwards of 25% of all BUSCO genes, while RE-associated BUSCOs averaged ∼1-2% of all BUSCOs in Hymenoptera and Diptera. To address whether general trends in RE-associated BUSCOs could be driven by an artifact of assembly errors (which might simply be more numerous in larger assemblies), rather than true associations between REs and BUSCO genes, we predicted that less contiguous, short-read assemblies would show inflated RE-BUSCO associations compared to more contiguous, long-read assemblies. However, this comparison revealed the opposite pattern; RE-associated BUSCOs are ∼60% more common in long-read assemblies (Welch two-sample *t*-test, *P* = 0.007; Supplemental Fig. S7).

Since most RE annotation relies on reference databases (i.e., Repbase, Dfam, (Hubley et al. 2016; Jurka et al. 2005), we expected bias in database representation to impact our RE annotation. The proportion of unclassified REs in a given assembly increased with its genetic distance from *D. melanogaster* (Spearman’s correlation = 0.4, *P* < 2 x 10^16^; Fig. 3a). For reference, unclassified repeats comprised only 13.1% of all repeats in the 71 *Drosophila* species but accounted for 40.5% total repeats on average in all other taxa. High fractions of unclassified repeats were especially evident in poorly sampled, early diverging insect orders. For example, in Thysanoptera and Ephemeroptera, 72.0% and 85.1% of respective REs are unclassified despite having similar genomic proportions of REs as *Drosophila* (∼25% in all three groups; Fig. 3b). Unclassified repeats were typically short sequences (mean length = 188.9 bp) and were slightly longer in long-read compared to short-read assemblies (Supplemental Fig. S8).

**Figure 3.**
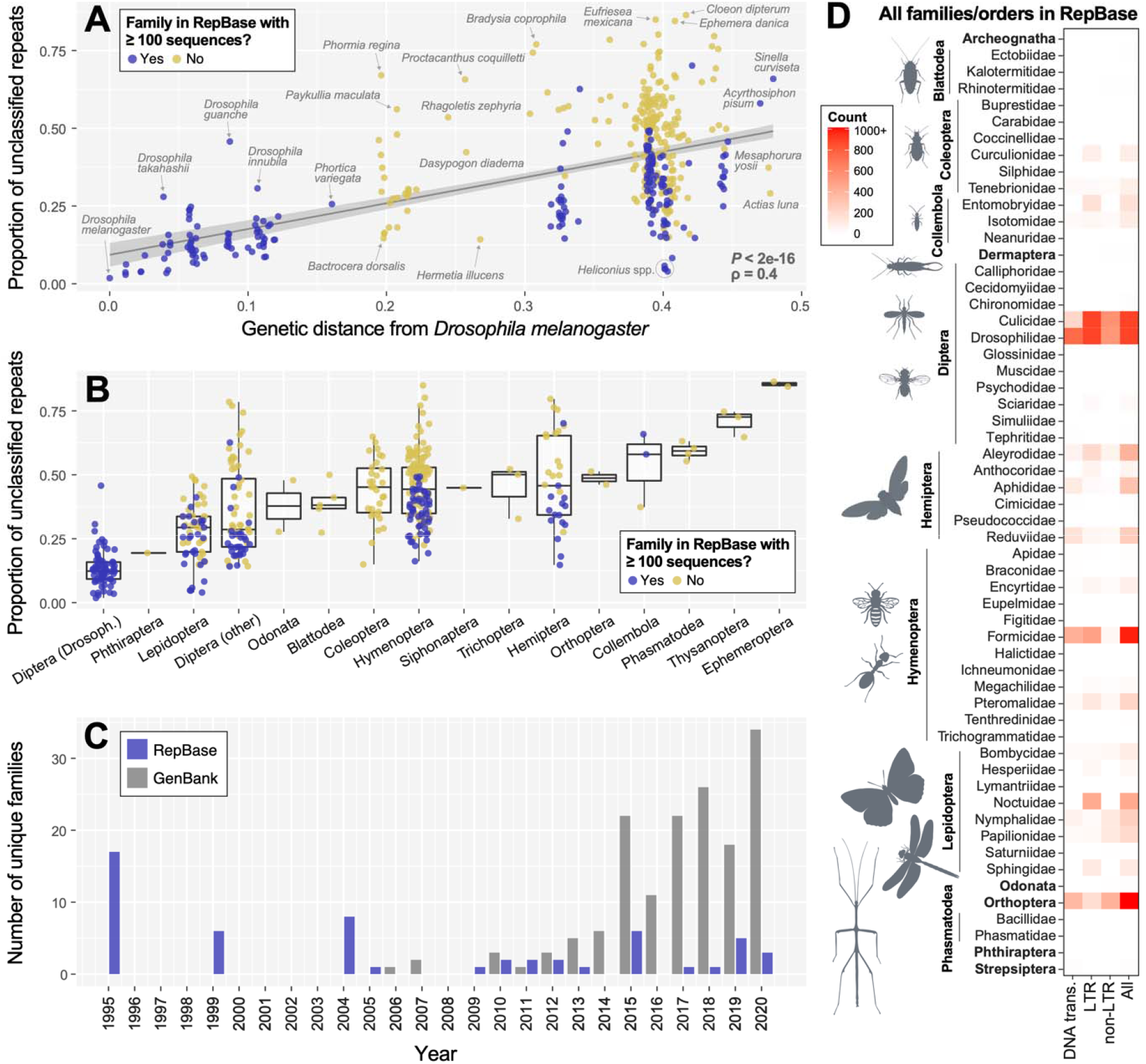
Insect representation in repetitive element databases and effects on RE detection. (A) A comparison of the proportion of total repeats that are unclassified in each insect’s genome assembly versus its genetic distance from *Drosophila melanogaster*. (B) The same data presented in (A) but grouped by order except for Diptera which are divided into family Drosophilidae and all other Diptera. In both (A) and (B), a “yes” reflects insect family-level representation of 100 or more sequences in Repbase. (C) Unique entries at the insect family-level submitted to Repbase or GenBank from 1995–2020. Data for GenBank submissions were taken from Hotaling et al. (2021). Of note, for 2020, only GenBank submissions through October 2020 were included. (D) Heatmap showing the abundance (count) of RE sequence entries in Repbase by order (bold) or family. Of the 154 insect families in our dataset, roughly one third, those listed here, have any representation in Repbase. Of those, many are represented by few RE sequences, e.g., essentially white boxes indicate only 1–10 sequences are present. If a single insect family was present, it is labeled with the broader order name, if two or more insect families from the same order were present, they are listed with a line encompassing them to the left.

To clarify the impact of uneven taxonomic representation in reference databases on RE annotation, we quantified representation of insect orders and families in Repbase (Bao et al. 2015; Jurka et al. 2005). Repbase is the most widely cited repository of RE sequences and is integral to the standard RE identification and annotation programs RepeatModeler2 (Flynn et al. 2020) and RepeatMasker (Smit and Hubley). Of the 20 insect orders in our data set, 14 are represented in Repbase, however, of those six are represented by a single family (Fig. 3d, Table S2). Of 154 insect families in our data set, just over one-third (*n* = 57) had any representation in Repbase.

Taxonomic bias is more extreme when the number of reference sequences is considered. Just two families, Drosophilidae and Culicidae (order Diptera), account for ∼60% (*n* = 8,453) of all insect sequences in Repbase (Fig. 3d) and ∼70% of all LTR sequences (*n* = 5,908). Nearly 75% of all insect families in Repbase are represented by fewer than 100 sequences and only four families [Culicidae (Diptera), Drosophilidae (Diptera), Formicidae (Hymenoptera), and Acrididae (Orthoptera)] have > 1000 sequences (Fig. 3d, Table S3). Species belonging to a family represented by ≥ 100 sequences in Repbase had, on average, 24.5% unclassified REs, whereas insects belonging to families represented by ≤ 99 sequences had nearly double the proportion of unclassified repeats (45.8%). The gap between available genome assemblies for insects and Repbase representation appears to be increasing. Since insect genome assemblies began to proliferate on GenBank in ∼2010, submissions to Repbase have not exhibited similar growth (Fig. 3c).

## Discussion

In the present study, we extended previous efforts (e.g., Gilbert et al. 2021; Petersen et al. 2019) by describing RE dynamics for 600+ insect species. In the process, we evaluated the efficacy of automated RE annotation pipelines in a large, taxonomically diverse data set and clarified expectations for RE annotation success in diverse clades.

### REs in insects: new insight from a broad taxonomic comparison

Insects account for over half of all described animal species (Stork 2018). To understand the genomic basis of this diversity, we must understand repeat evolution, as repeats comprise major fractions of nearly all insect genomes. We identified wide variation in RE abundance both within and among major clades. For example, DNA transposons were generally abundant in most insect orders yet conspicuously uncommon in Lepidoptera. Similarly, LINEs are abundant in many orders (e.g., Coleoptera, Trichoptera, Hemiptera) but largely absent in Hymenoptera (Figs. 1–2). These order-level patterns indicate both deep phylogenetic constraints in RE architecture (e.g., within orders), as well as major shifts between lineages. For example, in Holometabola, LINEs shift from low abundance in Hymenoptera to higher abundance in the next-branching lineages (i.e., Lepidoptera, Trichoptera, Coleoptera), then back to lower abundance in Diptera (Fig. 2m), suggesting shifts in strategies for maintaining genome stability and TE regulation across groups.

Our analysis of RE-associated BUSCOs illustrates how the evolution of interspersed repeats within and around protein-coding genes has evolved dynamically across lineages (Figs. 1C and 2m). Given the strong correlation between LINE abundance and the presence of RE-associated BUSCOS across insects (Supplemental Fig. S6), lineages with abundant LINEs may experience elevated rates of evolution in genomic regions, with potentially broad consequences for phenotype evolution. Genomes suppress RE activity through epigenetic silencing of repetitive sequences (e.g., heterochromatin formation, Slotkin and Martienssen 2007). Because silent marks may occur near regulatory gene regions and spread to adjacent sequences (Lee and Karpen 2017; Wei et al. 2022), movement of REs to new genomic loci can have immediate impact on expression of nearby genes. Over longer timescales, RE sequences can be co-opted to form genome-wide regulatory networks of gene expression (Chuong et al. 2017). Although we don’t present direct evidence of phenotype impacts here, our finding of abundant and dynamically evolving RE-gene associations in insects identifies new potential for studying RE impacts on coding regions and phenotype evolution in insects.

Our broad taxonomic sampling illustrates that non-model insects tend to have larger, more repeat-rich genomes than the model species (e.g., *D. melanogaster*) that seeded much of our present knowledge of RE dynamics (Fig. 1a–b, see also Supplemental Results). While REs can have both deleterious and adaptive impacts on host genomes (González et al. 2008; Petrov 2002), their dynamics are understudied in complex, repeat-rich genomes. The few larger-genome model groups (e.g. >1000 Mb) that have been comprehensively studied for REs (e.g., maize) suggest an ecosystem-like environment where REs adopt diverse strategies and impacts within their various niches in the genome (Stitzer et al. 2021). Investigating the diversity of insect models with varying genome sizes and complexity can expand our perspectives on genome evolution. For example, in caddisflies (Trichoptera), clades containing relatively larger genomes (e.g., 600– 2100 Mb) show higher species diversity and ecological breadth than small-genome lineages (e.g., >600 Mb), raising the potential for adaptive advantages of maintaining high repeat loads (Heckenhauer et al. 2022; Olsen et al. 2021). While the current study includes assembly lengths up to 4100 Mb, flow cytometry data shows evidence of insect genomes exceeding 18,000 Mb (*Bryodemella holdereri*, Cong et al. 2022). Insect models have potential to offer new insights on genome gigantism as assemblies for additional large-genome species become available (Liu et al. 2022). With high species diversity and broad distributions in nearly all habitat types, insects may be particularly useful for understanding factors driving temporal dynamics of TE activity including population demographics (Schrader et al. 2014) and environmental stress (Horváth et al. 2017; Signor et al. 2022). In addition, a “many-model” phylogenetic framework offered by insects may be key to connecting patterns of genome size, REs, and developmental constraints with ecological factors (Blommaert 2020).

### Sequencing technology and RE analysis

Our analyses showed that sequencing technology influences RE detection. Specifically, long-read assemblies contain 36% more REs than short-read assemblies. LINEs and DNA transposons showed low or modest impact from technology differences (e.g., differences in LINEs detection were non-significant, Fig. 2g). This, combined with their overall genomic abundance, even in lineages with poor representation in RE databases (e.g., Coleoptera, Blattodea; Figs. 1, 3d), suggests robustness to both technology differences and limited database representation for LINEs and DNA transposons. By contrast, LTRs showed 162% increase in long-read assemblies. LTRs are difficult to identify with standard approaches due to their length and sequence complexity (Flynn et al. 2020) and this finding suggests technology advances are closing the assembly and annotation gaps for historically problematic elements. Other recent studies that report telomere-to-telomere assemblies (e.g., Miga et al. 2020) and improved contiguity through combining data from multiple sequencing strategies (e.g., Peona et al. 2021) further illustrate the impact of technology advances in resolving assemblies at repetitive regions.

Tandem repeats may now be the last RE type for which assembly remains largely intractable. Although long-read assemblies showed modest gains in tandem repeat detection (∼25% increase), large blocks (e.g., megabases) of tandem repeats such as satellite DNAs are common in insects and other groups (UgarkoviĆ and Plohl 2002) and will remain unresolved in assemblies for the near future. Assembly-free approaches that estimate RE abundance from raw reads (e.g, through clustering algorithms like RepeatExplorer2, Nóvak et al. 2010) remain important tools for estimating abundance of repeats that may be collapsed in assemblies -- especially tandem repeats or TEs with a recent history of expansion. Indeed, our comparison of RE estimates between long-read assemblies and cluster-based methods corroborate the importance of these programs, as our cluster-based analysis consistently showed increased detection of tandem repeats over the long-read assembly estimates (Supplemental Figs. S4 and S5). Our analysis offers an additional insight that clustering-based programs may also be prone to large underestimates of interspersed repeats that appear to be much better detected by assembly-based approaches. These trends illustrate how the repeat architecture of specific repeat categories within a given genome are likely to impact which analysis approach is most effective and reinforce the importance of applying multiple orthogonal methods.

Our findings complement studies that report within-species comparisons of the impact of sequencing technology (Solares et al. 2018; Rech et al. 2022) on TE detection by leveraging insect diversity to provide a broad perspective informed by hundreds of species. In addition to general trends in long-vs short-read assemblies reported here, finer layers of technology-related factors such as the impact of specific sequencing platforms (e.g., PacBio CLR vs PacBio HiFi) (Chu et al. 2021), genome assembly algorithms (Chang et al. 2019), and genome assembly/TE detection tools (Bergman and Quesneville 2007; Goerner-Potvin and Bourque 2018) are expected to impact TE detection on a per-assembly basis. Ongoing studies designed to identify technical protocols that maximize resolution of TEs in assemblies are needed to guide best practices in the face of constantly changing sequencing and assembly technology.

### Challenges and opportunities for RE biology in biodiversity genomics

We provide an empirical illustration of fundamental challenges that limit thorough RE annotation in all but a few model species and their close relatives. Given the scale of repeats that could not be annotated in many lineages (i.e., unclassified repeats, Figs. 1, 3a–b) we show how deep insights into RE dynamics across phylogenetic scales remain impractical until we can map the finer details of RE landscapes in any species.

To realize the potential that biodiversity genomics offers for the study of REs in insects and beyond, we must be able to efficiently study homologous REs across clades. Two main challenges have slowed progress toward this goal: assembly fragmentation in repetitive regions and comprehensive RE annotation. The rise of long-read sequencing technology has improved assembly of repetitive regions (e.g., Hotaling et al. 2021b) and largely ameliorated this first challenge. This advance has been driven primarily by industry research and incentives paired with buy-in from the genomics community, including consortia (e.g., Earth BioGenome Project, (Lewin et al. 2018). However, advances in RE annotation rely largely on the academic community with fewer financial or related incentives. Although many tools for automated identification and annotation of REs exist (Bergman and Quesneville 2007; Goerner-Potvin and Bourque 2018), annotation tools are limited by the quality of reference databases and specifically the breadth of known REs that can be used to annotate unknown REs in focal assemblies.

As such, community led RE database curation is not trivial. Two specific obstacles to effective annotation exist: (**1**) RE taxonomies are in early stages of curation such that redundantly described, or undescribed REs are both common; and (**2**) taxonomic representation in existing RE databases is woefully incomplete (Fig. 3d). Although these issues have been raised in the RE community for more than a decade (Bergman and Quesneville 2007; Elliott et al. 2021; Hoen et al. 2015; Piégu et al. 2015), our results add quantification to an abstract challenge and highlight that despite major progress in biodiversity genomics overall, the RE “database issue” is growing worse rather than improving (Fig. 3c). As it stands, an average of 40.5% of total repeats could not be classified with our automated classification approach in all non-*Drosophila* taxa, while just 13.1% are unclassified on average in the 71 *Drosophila* species sampled (Fig. 3b). The numbers are much worse in early diverging insect orders such as Thysanoptera and Ephemeroptera (72.0% and 85.1% unclassified, respectively) despite their having similar genomic proportions of REs as *Drosophila* (∼25% in all three groups; Fig. 3b). Without a concerted effort to improve RE curation and annotation, we expect unclassified percentages of REs to increase as additional assemblies are sequenced from new species. These problems are not likely specific to insects and present a fundamental challenge that impedes deep understanding of genomes that genomicists seek.

To be clear, we applaud the efforts of many groups that develop, maintain, and curate RE repositories such as Repbase and Dfam (Bao et al. 2015; Hubley et al. 2016; Jurka et al. 2005; Storer et al. 2021; Wheeler et al. 2012). We also acknowledge the valuable efforts from research groups studying classical model species (e.g., *Drosophila melanogaster*) whose contributions form a basis of broad understanding about RE biology. As biodiversity genomics continues to grow and diversify, concerted efforts should be made to support RE research and make the importance of their annotation central to broader goals of the field (i.e., similar to generating new genome assemblies or gene annotation tools).

We view biodiversity science as a large-scale solution to many challenges facing RE biology. With a long history of deep expertise in phylogenetics, taxonomy, and specimen acquisition, the infrastructure, experience, and human resources within biodiversity science could be a boon for improving RE taxonomy, curation, and taxonomic representation. However, we emphasize the need for care when embracing this challenge. A primary lesson learned from decades of taxonomy and phylogenetic inference is that thorough taxon sampling is critical to avoid mistakes in both endeavors. Thus, a stable RE taxonomy hinges upon the mapping of REs in taxonomically diverse groups, establishing homology through robust phylogenetic analysis of specific elements within and across groups, *and* submitting curated RE sequences to existing databases (Bao et al. 2015; Hubley et al. 2016; Jurka et al. 2005; Storer et al. 2021; Wheeler et al. 2012). In turn, studying REs in diverse clades can offer reciprocal benefits to biodiversity science in that repetitive elements are an underutilized source of signal that can add resolution to evolutionary studies (Dodsworth et al. 2015; Negm et al. 2021; Sproul et al. 2020).

As we move forward in this new era of biodiversity genomics, we need to simultaneously meet the challenge of studying RE dynamics across broad taxonomic scales. To bridge this gap, we offer three ways for the genomics community to contribute:

1. **Embrace RE biology.** Rather than viewing REs as nuisance sequences to be masked (Slotkin 2018), seek to understand their interesting and diverse roles in genome biology. Many excellent, accessible reviews exist (e.g., Bourque et al. 2018; Wells and Feschotte 2020) and more RE literacy and interest will no doubt improve RE science.
2. **Document REs in new (and existing) genome assemblies.** Whether generating a new genomic resource or evaluating one as a reviewer, editor, or peer, encourage reporting and documentation of REs. This will add to the RE knowledge base and accelerate literacy of both REs and the software tools available for their study.
3. **Invest in RE library curation and database submission within your area of taxonomic expertise.** To meet the challenge of RE annotation with accelerating availability of genome assemblies, RE library curation and database submission needs to become a mainstream step in data archiving. There are many resources designed to streamline contribution and data sharing, including RE curation guidelines (Goubert et al. 2022), descriptions of Repbase and Dfam databases and submission (Kohany et al. 2006; Storer et al. 2021), TE library curation tools (e.g., Ou et al. 2019), and group-specific RE resources (Elliott et al. 2021).

From single, difficult-to-obtain genome assemblies ∼20 years ago to dozens of new, highly contiguous assemblies being published every day, an exciting, new discipline of biodiversity genomics has emerged. By investing in solutions to address bottlenecks for studying REs and any similar challenges, we can build the foundation for unprecedented new understanding of genome biology in insects and across the tree of life.

## Methods

An extended version of Materials and Methods with additional details of phylogenetic inference, and RE-gene associations, and statistical analysis is provided in Supplemental Materials.

### Data acquisition

Following Hotaling et al. (2020), we used the assembly-descriptors function in the NCBI datasets command line tool to download metadata for all nuclear genomes available for insects on GenBank (accessed 2 November 2020; (Sayers et al. 2021). We then culled our data set to include only one representative genome per taxon (species or subspecies) by selecting the assembly with the highest contig N50 (the mid-point of the contig distribution where 50% of the genome is assembled into contigs of a given length or longer). Using provided NCBI metadata on the sequencing read technology used for assembly, assemblies were classified as “short-read”, “long-read”, or “not provided” based on whether only short-reads (e.g., Illumina) were used, any amount of long-read sequences (e.g., PacBio) were used, or no information was provided. After identifying our focal genome set, we downloaded the relevant genomes for downstream analysis. Analysis scripts used in this study, including those that were used for data collection, are included in this study’s GitHub repository. A full list of the genome assemblies used in this study are provided in Supplemental Table S1 (Supplemental Material).

### Quantifying assembly completeness and phylogenetic inference

To assess gene completeness, we ran “Benchmarking Universal Single Copy Orthologs” (BUSCO) v.4.1.4 (Seppey et al. 2019) on each assembly using the 1,367 reference genes in the OrthoDB v.10 Insecta gene set (Kriventseva et al. 2019) and the “--long” analysis mode. We divided our data set into three subsets: (1) the full data set with no filtering, (2) only assemblies with BUSCO gene content ≥ 50%, or (3) only assemblies with BUSCO gene content ≥ 90%. To organize our results in a phylogenetic framework, we then estimated a species tree for our full data set using single-copy orthologs resulting from the BUSCO analyses.

### Repeat element identification and annotation

We identified REs in genome assemblies using RepeatModeler2.0 (Flynn et al. 2020) with search engine “ncbi”, which also generates a library of repeat consensus sequences. We annotated repeats in assemblies through two rounds of annotation with RepeatMasker4.1.0 (Smit and Hubley), the first round used custom repeat libraries generated by RepeatModeler2 for each respective assembly and with search engine “ncbi” and option -xsmall. We then converted the softmasked assembly resulting from the first RepeatMasker round to a hardmasked assembly using the lc2n.py script (https://github.com/PdomGenomeProject/repeat-masking), and re-ran RepeatMasker on the hard-masked assembly with RepeatMasker’s internal arthropod repeat library and species “Arthropoda”. RepeatMasker’s internal library, RepeatMaskerLib.embl combines elements from Repbase, Dfam, and Artefacts data repositories per software documentation. (Based on our analysis of repository composition, the large majority of insect models in public repositiories at the date of this research are in Repbase.) We then merged RepeatMasker output tables from both runs to summarize the abundance of RE categories. We studied patterns of repeat dynamics within and across taxonomic groups by parsing RepeatMasker output tables and visualizing the distribution and abundance of major RE categories using custom Python and R scripts.

As an orthogonal approach to identifying repetitive elements with our assembly-based analysis, we explored genome repetitiveness and RE abundance with assembly-free approaches based on clustering of low-coverage short-read data as implemented in both dnaPipeTE v1.3.1 (Goubert et al. 2015) and RepeatExplorer2 (Novák et al. 2013). The former program relies on similar dependencies (e.g., RepeatMasker and Dfam and Repbase repeat databases) as our assembly-based approach and is thus well suited to exploring the effects of a clustering-based approach on RE estimates, while reducing potential noise introduced by program-specific software and database dependencies. RepeatExplorer2 provides a reference point for a similar tool that uses a different underlying repeat database (i.e., Metazoa 3.0) and dependencies, including TAREAN (Novák et al. 2017) which specializes in identification of satellite DNAs. Samples for these analyses were chosen both to spread taxonomic representation across multiple insect orders, and to minimize potential noise introduced by variation in technical details surrounding data generation. Additional details for clustering-based analyses are provided in Supplemental Materials.

### Correlation analyses

We tested for correlations between RE abundance and a range of aspects for each genome assembly, including sequencing technology, using R version 3.6.3 (R Core Team 2013). These included a comparison of total REs identified as well as specific classes (e.g., LINEs) versus the primary sequencing technology used (short- or long-reads). For all correlation analyses, we tested for normality in our data sets with a Shapiro-Wilk test and since the null hypothesis was rejected for all data sets (*P* < 0.05), we used Spearman’s rank correlation tests.

### Repetitive element and protein-coding gene associations

For all assemblies with ≥ 90% BUSCO gene content, we measured RE-gene associations (i.e., RE sequences inserted within or adjacent to protein-coding genes) following Heckenhauer et al. (Heckenhauer et al. 2022). Their study validated a new approach to quantifying RE sequences associated with BUSCOs. In some cases, RE fragments are embedded within BUSCOs, and in others, REs with open reading frames that are immediately adjacent to BUSCOs are inadvertently classified by the BUSCO algorithm as being part of the BUSCO. They showed that quantifying such instances of RE sequences in BUSCOs can serve as a proxy for genome-wide RE-gene associations. Our approach adapted theirs to suit a higher throughput analysis, as described in more detail in Supplemental Materials.

### Investigating the effects of taxonomic sampling bias

We investigated effects of taxonomic sampling bias on our understanding of REs in insects by analyzing the composition of the Repbase repository for RE sequences and the resulting impact on repeat annotation in our assemblies. We used custom scripts to parse the Repbase database and quantify the taxonomic representation of insect orders and families included in our data set, as well as the rate of insect repeat submissions over time.

## Data access

Species-specific repeat libraries generated by RepeatModeler2 and summary tables are available on FigShare (https://doi.org/10.6084/m9.figshare.c.6024905.v1) and Supplemental Materials and have been submitted to Dfam (accessions: DR2407971–DR3440067). All scripts used in analyses are available on GitHub (https://github.com/johnssproul/Insect_REs) and Supplemental Materials.

## Acknowledgments

We are thankful to Robert Hubley and Jessica Storer for their support preparing our species-specific RepeatModeler libraries upload to the Dfam database. JSS was supported by a National Science Foundation Postdoctoral Research Fellowship in Biology (Division of Biological Infrastructure DBI-1811930) and a National Institutes of Health General Medical Sciences grant (R35GM119515) awarded to AML. SH and JLK were supported by NSF award #OPP-1906015. JH is member of the DFG funded priority programme, Genomic Basis of Evolutionary Innovations (GEvol): SPP2349 funded by the Deutsche Forschungsgemeinschaft (DFG, German Research Foundation), project number: 502865717. J.H and S.U.P. were supported by LOEWE-Centre for Translational Biodiversity Genomics funded by the Hessen State Ministry of Higher Education, Research and the Arts (HMWK).

## References

Bao W, Kojima KK, Kohany O. 2015. Repbase Update, a database of repetitive elements in eukaryotic genomes. Mob DNA 6: 11.

Bennetzen JL. 2000. Transposable element contributions to plant gene and genome evolution. Plant Mol Biol 42: 251–269.

Bergman CM, Quesneville H. 2007. Discovering and detecting transposable elements in genome sequences. Brief Bioinform 8: 382–392.

Blommaert J. 2020. Genome size evolution: towards new model systems for old questions. Proc R Soc B Biol Sci 287: 20201441.

Bourque G, Burns KH, Gehring M, Gorbunova V, Seluanov A, Hammell M, Imbeault M, Izsvák Z, Levin HL, Macfarlan TS, et al. 2018. Ten things you should know about transposable elements. Genome Biol 19: 199.

Brand CL, Levine MT. 2022. Cross-species incompatibility between a DNA satellite and the Drosophila Spartan homolog poisons germline genome integrity. Curr Biol. https://www.sciencedirect.com/science/article/pii/S0960982222007680 (Accessed May 31, 2022).

Cáceres M, Ranz JM, Barbadilla A, Long M, Ruiz A. 1999. Generation of a Widespread Drosophila Inversion by a Transposable Element. Science 285: 415–418.

Chang C-H, Larracuente AM. 2019. Heterochromatin-Enriched Assemblies Reveal the Sequence and Organization of the Drosophila melanogaster Y Chromosome. Genetics 211: 333–348.

Chu C, Zhao B, Park PJ, Lee EA. 2020. Identification and Genotyping of Transposable Element Insertions From Genome Sequencing Data. Curr Protoc Hum Genet 107: e102.

Chuong EB, Elde NC, Feschotte C. 2017. Regulatory activities of transposable elements: from conflicts to benefits. Nat Rev Genet 18: 71–86.

Cong Y, Ye X, Mei Y, He K, Li F. 2022. Transposons and non-coding regions drive the intrafamily differences of genome size in insects. iScience 25: 104873.

de Koning AJ, Gu W, Castoe TA, Batzer MA, Pollock DD. 2011. Repetitive elements may comprise over two-thirds of the human genome. PLoS Genet 7: e1002384.

Dodsworth S, Chase MW, Kelly LJ, Leitch IJ, Macas J, Novák P, Piednoël M, Weiss-Schneeweiss H, Leitch AR. 2015. Genomic Repeat Abundances Contain Phylogenetic Signal. Syst Biol 64: 112–126.

Elliott TA, Heitkam T, Hubley R, Quesneville H, Suh A, Wheeler TJ, The TE Hub Consortium. 2021. TE Hub: A community-oriented space for sharing and connecting tools, data, resources, and methods for transposable element annotation. Mob DNA 12: 16.

Ferree PM, Barbash DA. 2009. Species-specific heterochromatin prevents mitotic chromosome segregation to cause hybrid lethality in Drosophila. PLOS Biol 7: e1000234.

Flynn JM, Hubley R, Goubert C, Rosen J, Clark AG, Feschotte C, Smit AF. 2020. RepeatModeler2 for automated genomic discovery of transposable element families. Proc Natl Acad Sci 117: 9451–9457.

Gilbert C, Peccoud J, Cordaux R. 2021. Transposable Elements and the Evolution of Insects. Annu Rev Entomol 66: 355–372.

Goerner-Potvin P, Bourque G. 2018. Computational tools to unmask transposable elements. Nat Rev Genet 1.

González J, Lenkov K, Lipatov M, Macpherson JM, Petrov DA. 2008. High Rate of Recent Transposable Element–Induced Adaptation in Drosophila melanogaster. PLOS Biol 6: e251.

Goubert C, Craig RJ, Bilat AF, Peona V, Vogan AA, Protasio AV. 2022. A beginner’s guide to manual curation of transposable elements. Mob DNA 13: 7.

Goubert C, Modolo L, Vieira C, ValienteMoro C, Mavingui P, Boulesteix M. 2015. De Novo Assembly and Annotation of the Asian Tiger Mosquito (Aedes albopictus) Repeatome with dnaPipeTE from Raw Genomic Reads and Comparative Analysis with the Yellow Fever Mosquito (Aedes aegypti). Genome Biol Evol 7: 1192–1205.

Heckenhauer J, Frandsen PB, Sproul JS, Li Z, Paule J, Larracuente AM, Maughan PJ, Barker MS, Schneider JV, Stewart RJ, et al. 2022. Genome size evolution in the diverse insect order Trichoptera. GigaScience 11: giac011.

Hoen DR, Hickey G, Bourque G, Casacuberta J, Cordaux R, Feschotte C, Fiston-Lavier A-S, Hua-Van A, Hubley R, Kapusta A, et al. 2015. A call for benchmarking transposable element annotation methods. Mob DNA 6: 13.

Horváth V, Merenciano M, González J. 2017. Revisiting the Relationship between Transposable Elements and the Eukaryotic Stress Response. Trends Genet 33: 832–841.

Hotaling S, Kelley JL, Frandsen PB. 2020. Aquatic Insects Are Dramatically Underrepresented in Genomic Research. Insects 11: 601.

Hotaling S, Kelley JL, Frandsen PB. 2021a. Toward a genome sequence for every animal: Where are we now? Proc Natl Acad Sci 118: e2109019118.

Hotaling S, Sproul JS, Heckenhauer J, Powell A, Larracuente AM, Pauls SU, Kelley JL, Frandsen PB. 2021b. Long Reads Are Revolutionizing 20 Years of Insect Genome Sequencing. Genome Biol Evol 13: evab138.

Hubley R, Finn RD, Clements J, Eddy SR, Jones TA, Bao W, Smit AF, Wheeler TJ. 2016. The Dfam database of repetitive DNA families. Nucleic Acids Res 44: D81–D89.

Jagannathan M, Yamashita YM. 2021. Defective satellite DNA clustering into chromocenters underlies hybrid incompatibility in Drosophila. Mol Biol Evol 38: 4977–4986.

Jurka J, Kapitonov VV, Pavlicek A, Klonowski P, Kohany O, Walichiewicz J. 2005. Repbase Update, a database of eukaryotic repetitive elements. Cytogenet Genome Res 110: 462–467.

Kim YB, Oh JH, McIver LJ, Rashkovetsky E, Michalak K, Garner HR, Kang L, Nevo E, Korol AB, Michalak P. 2014. Divergence of Drosophila melanogaster repeatomes in response to a sharp microclimate contrast in Evolution Canyon, Israel. Proc Natl Acad Sci 111: 10630–10635.

Kohany O, Gentles AJ, Hankus L, Jurka J. 2006. Annotation, submission and screening of repetitive elements in Repbase: RepbaseSubmitter and Censor. BMC Bioinformatics 7: 474.

Kriventseva EV, Kuznetsov D, Tegenfeldt F, Manni M, Dias R, Simão FA, Zdobnov EM. 2019. OrthoDB v10: sampling the diversity of animal, plant, fungal, protist, bacterial and viral genomes for evolutionary and functional annotations of orthologs. Nucleic Acids Res 47: D807–D811.

Lee YCG, Karpen GH. 2017. Pervasive epigenetic effects of Drosophila euchromatic transposable elements impact their evolution ed. M. Nordborg. eLife 6: e25762.

Lewin HA, Robinson GE, Kress WJ, Baker WJ, Coddington J, Crandall KA, Durbin R, Edwards SV, Forest F, Gilbert MTP, et al. 2018. Earth BioGenome Project: Sequencing life for the future of life. Proc Natl Acad Sci 115: 4325–4333.

Liu X, Majid M, Yuan H, Chang H, Zhao L, Nie Y, He L, Liu X, He X, Huang Y. 2022. Transposable element expansion and low-level piRNA silencing in grasshoppers may cause genome gigantism. BMC Biol 20: 243.

Marks RA, Hotaling S, Frandsen PB, VanBuren R. 2021. Representation and participation across 20 years of plant genome sequencing. Nat Plants 7: 1571–1578.

Miga KH, Koren S, Rhie A, Vollger MR, Gershman A, Bzikadze A, Brooks S, Howe E, Porubsky D, Logsdon GA, et al. 2020. Telomere-to-telomere assembly of a complete human X chromosome. Nature 585: 79–84.

Misof B, Liu S, Meusemann K, Peters RS, Donath A, Mayer C, Frandsen PB, Ware J, Flouri T, Beutel RG, et al. 2014. Phylogenomics resolves the timing and pattern of insect evolution. Science 346: 763–767.

Negm S, Greenberg A, Larracuente AM, Sproul JS. 2021. RepeatProfiler: a pipeline for visualization and comparative analysis of repetitive DNA profiles. Mol Ecol Resour 21: 969–981.

>Novák P, Ávila Robledillo L, KoblíŽková A, Vrbová I, Neumann P, Macas J. 2017. TAREAN: a computational tool for identification and characterization of satellite DNA from unassembled short reads. Nucleic Acids Res 45: e111–e111.

Novák P, Guignard MS, Neumann P, Kelly LJ, Mlinarec J, KoblíŽková A, Dodsworth S, Kovařík A, Pellicer J, Wang W, et al. 2020. Repeat-sequence turnover shifts fundamentally in species with large genomes. Nat Plants 6: 1325–1329.

Novák P, Neumann P, Pech J, Steinhaisl J, Macas J. 2013. RepeatExplorer: a Galaxy-based web server for genome-wide characterization of eukaryotic repetitive elements from next-generation sequence reads. Bioinformatics 29: 792–793.

Novák P, Neumann P, Macas J. 2010. Graph-based clustering and characterization of repetitive sequences in next-generation sequencing data. BMC Bioinformatics 11: 378.

Olsen LK, Heckenhauer J, Sproul JS, Dikow RB, Gonzalez VL, Kweskin MP, Taylor AM, Wilson SB, Stewart RJ, Zhou X, et al. 2021. Draft Genome Assemblies and Annotations of Agrypnia vestita Walker, and Hesperophylax magnus Banks Reveal Substantial Repetitive Element Expansion in Tube Case-Making Caddisflies (Insecta: Trichoptera). Genome Biol Evol 13: evab013.

Ou S, Su W, Liao Y, Chougule K, Agda JRA, Hellinga AJ, Lugo CSB, Elliott TA, Ware D, Peterson T, et al. 2019. Benchmarking transposable element annotation methods for creation of a streamlined, comprehensive pipeline. Genome Biol 20: 275.

Peona V, Blom MPK, Xu L, Burri R, Sullivan S, Bunikis I, Liachko I, Haryoko T, Jønsson KA, Zhou Q, et al. 2021. Identifying the causes and consequences of assembly gaps using a multiplatform genome assembly of a bird-of-paradise. Mol Ecol Resour 21: 263–286.

Petersen M, Armisén D, Gibbs RA, Hering L, Khila A, Mayer G, Richards S, Niehuis O, Misof B. 2019. Diversity and evolution of the transposable element repertoire in arthropods with particular reference to insects. BMC Ecol Evol 19: 11.

Petrov DA. 2002. Mutational Equilibrium Model of Genome Size Evolution. Theor Popul Biol 61: 531–544.

Piégu B, Bire S, Arensburger P, Bigot Y. 2015. A survey of transposable element classification systems–a call for a fundamental update to meet the challenge of their diversity and complexity. Mol Phylogenet Evol 86: 90–109.

R Core Team. 2013. R: A language and environment for statistical computing. R Found Stat Comput Vienna Austria URL Http://www.R-Proj.

Rech GE, Radío S, Guirao-Rico S, Aguilera L, Horvath V, Green L, Lindstadt H, Jamilloux V, Quesneville H, González J. 2022. Population-scale long-read sequencing uncovers transposable elements associated with gene expression variation and adaptive signatures in Drosophila. Nat Commun 13: 1–16.

Sayers EW, Cavanaugh M, Clark K, Pruitt KD, Schoch CL, Sherry ST, Karsch-Mizrachi I. 2021. GenBank. Nucleic Acids Res 49: D92–D96.

Schley RJ, Pellicer J, Ge X-J, Barrett C, Bellot S, Guignard MS, Novák P, Suda J, Fraser D, Baker WJ, et al. 2022. The ecology of palm genomes: repeat-associated genome size expansion is constrained by aridity. New Phytologist 236: 433–446.

Schrader L, Kim JW, Ence D, Zimin A, Klein A, Wyschetzki K, Weichselgartner T, Kemena C, Stökl J, Schultner E, et al. 2014. Transposable element islands facilitate adaptation to novel environments in an invasive species. Nat Commun 5: 5495.

Schrader L, Schmitz J. 2019. The impact of transposable elements in adaptive evolution. Mol Ecol 28: 1537–1549.

Seppey M, Manni M, Zdobnov EM. 2019. BUSCO: Assessing Genome Assembly and Annotation Completeness. In Gene Prediction: Methods and Protocols (ed. M. Kollmar), Methods in Molecular Biology, pp. 227–245, Springer, New York, NY 10.1007/978-1-4939-9173-0_14 (Accessed June 1, 2022).

Signor S, Yocum G, Bowsher J. 2022. Life stage and the environment as effectors of transposable element activity in two bee species. J Insect Physiol 137: 104361.

Slotkin RK. 2018. The case for not masking away repetitive DNA. Mob DNA 9: 15.

Slotkin RK, Martienssen R. 2007. Transposable elements and the epigenetic regulation of the genome. Nat Rev Genet 8: 272–285.

Smit A, Hubley R. RepeatMasker Open-4.1. 2019.

Solares EA, Chakraborty M, Miller DE, Kalsow S, Hall K, Perera AG, Emerson JJ, Hawley RS. 2018. Rapid low-cost assembly of the Drosophila melanogaster reference genome using low-coverage, long-read sequencing. G3 Genes Genomes Genet 8: 3143–3154.

Sotero-Caio CG, Platt RN, Suh A, Ray DA. 2017. Evolution and diversity of transposable elements in vertebrate genomes. Genome Biol Evol 9: 161–177.

Sproul JS, Barton LM, Maddison DR. 2020. Repetitive DNA profiles Reveal Evidence of Rapid Genome Evolution and Reflect Species Boundaries in Ground Beetles. Syst Biol. https://academic.oup.com/sysbio/advance-article/doi/10.1093/sysbio/syaa030/5817835 (Accessed May 15, 2020).

Stitzer MC, Anderson SN, Springer NM, Ross-Ibarra J. 2021. The genomic ecosystem of transposable elements in maize. PLOS Genet 17: e1009768.

Storer J, Hubley R, Rosen J, Wheeler TJ, Smit AF. 2021. The Dfam community resource of transposable element families, sequence models, and genome annotations. Mob DNA 12: 2.

Stork NE. 2018. How many species of insects and other terrestrial arthropods are there on Earth? Annu Rev Entomol 63: 31–45.

Stuart T, Eichten SR, Cahn J, Karpievitch YV, Borevitz JO, Lister R. 2016. Population scale mapping of transposable element diversity reveals links to gene regulation and epigenomic variation ed. D. Zilberman. eLife 5: e20777.

Sun L, Jing Y, Liu X, Li Q, Xue Z, Cheng Z, Wang D, He H, Qian W. 2020. Heat stress-induced transposon activation correlates with 3D chromatin organization rearrangement in Arabidopsis. Nat Commun 11: 1886.

UgarkoviĆ Ðjica, Plohl M. 2002. Variation in satellite DNA profiles—causes and effects. EMBO J 21: 5955–5959.

Vargas-Chavez C, Pendy NML, Nsango SE, Aguilera L, Ayala D, González J. 2022. Transposable element variants and their potential adaptive impact in urban populations of the malaria vector Anopheles coluzzii. Genome Res 32: 189–202.

Wei KH-C, Mai D, Chatla K, Bachtrog D. 2022. Dynamics and Impacts of Transposable Element Proliferation in the Drosophila nasuta Species Group Radiation. Mol Biol Evol 39: msac080.

Wells JN, Feschotte C. 2020. A field guide to eukaryotic transposable elements. Annu Rev Genet 54: 539–561.

Wheeler TJ, Clements J, Eddy SR, Hubley R, Jones TA, Jurka J, Smit AF, Finn RD. 2012. Dfam: a database of repetitive DNA based on profile hidden Markov models. Nucleic Acids Res 41: D70–D82.

Winter DJ, Ganley AR, Young CA, Liachko I, Schardl CL, Dupont P-Y, Berry D, Ram A, Scott B, Cox MP. 2018. Repeat elements organise 3D genome structure and mediate transcription in the filamentous fungus Epichloë festucae. PLoS Genet 14: e1007467.

