## Supplemental Materials for "600+ insect genomes reveal repetitive element dynamics and highlight biodiversity-scale repeat annotation challenges"

**This SI file contains:**
Supplemental Methods
Supplemental Results
References
Supplemental Figures

### Supplemental Methods

*Data acquisition*

Following Hotaling et al. (2020), we used the assembly-descriptors function in the NCBI datasets command line tool to download metadata for all nuclear genomes available for insects on GenBank (accessed 2 November 2020; Sayers et al. 2020). We then culled our data set to include only one representative genome per taxon (species or subspecies) by selecting the assembly with the highest contig N50 (the mid-point of the contig distribution where 50% of the genome is assembled into contigs of a given length or longer). Using provided NCBI metadata on the sequencing read technology used for assembly, assemblies were classified as "short-read", "long-read", or "not provided" based on whether only short-reads (e.g., Illumina) were used, any amount of long-read sequences (e.g., PacBio) were used, or no information was provided. After identifying our focal genome set, we downloaded the relevant genomes for downstream analysis. Analysis scripts used in this study, including those that were used for data collection, are included in this study's GitHub (https://github.com/johnssproul/Insect_REs). A full list of the genome assemblies used in this study are provided in Table S1.

*Quantifying assembly completeness and phylogenetic reconstruction*

To assess gene completeness, we ran "Benchmarking Universal Single Copy Orthologs" (BUSCO) v.4.1.4 (Seppey, et al. 2019) on each assembly using the 1,367 reference genes in the OrthoDB v.10 Insecta gene set (Kriventseva, et al. 2019) and the "--long" analysis mode. We divided our data set into three subsets: (1) the full data set with no filtering, (2) only assemblies with BUSCO gene content ≥ 50%, or (3) only assemblies with BUSCO gene content ≥ 90%. To organize our results in a phylogenetic framework, we then estimated a species tree for our full data set using single-copy orthologs resulting from the BUSCO analyses. We generated an unaligned FASTA file with sequences from each species for each single-copy ortholog. We then aligned each ortholog with the MAFFT L-INS-i algorithm (Katoh & Standley, 2013). We selected the best-fit substitution model for each alignment using ModelFinder (option -m mfp, (Kalyaanamoorthy et al., 2017) in IQtree v.2.0.6 (Minh et al., 2020) and estimated a maximum likelihood tree with 1000 ultrafast bootstrap replicates (Hoang et al., 2018) with BNNI correction (option -bb 1000 -bnni). We then generated a multi-species coalescent tree in ASTRAL-III (Zhang et al., 2018) using the best maximum likelihood tree from each ortholog as input. Because taxon sampling was not evenly distributed, some deep relationships within the tree conflicted with well-supported hypotheses from previous studies. To correct this, we modified the tree such that the ordinal-level relationships reflected those from Misof et al. (2014).

We generated a genetic distance matrix on a concatenated alignment of single copy orthologs to evaluate the effect of genetic distance from model organisms on the successful classification of REs. To do this, we first trimmed the single gene alignments using Aliscore v.0.2.2 (Misof and Misof 2009; Kück et al. 2010) and ALICUT v.2.31 (available from github: https://github.com/PatrickKueck/AliCUT). We then concatenated the ortholog alignments into a supermatrix using FASconCAT (Kück & Meusemann 2010). We estimated uncorrected pairwise distances in PAUP* v.4b (Swofford 2003) by adding the following nexus block to the supermatrix nexus file: "Begin paup;dset distance=p;savedist format=onecolumn;end;Quit;".

*Repeat element identification and annotation*
We identified REs in genome assemblies using RepeatModeler2.0 (Flynn et al. 2020) with search engine "ncbi", which also generates a library of repeat consensus sequences. We annotated repeats in assemblies through two rounds of annotation with RepeatMasker4.1.0 (Smit et al. 2020), the first round used custom repeat libraries generated by RepeatModeler2 for each respective assembly and with search engine "ncbi" and option -xsmall. We then the softmasked assembly resulting from the first RepeatMasker round to a hardmasked assembly using the lc2n.py script (https://github.com/PdomGenomeProject/repeat-masking), and re-ran RepeatMasker on the hard-masked assembly with RepeatMasker's internal arthropod repeat library and species "Arthropoda". We then merged RepeatMasker output tables from both runs to summarize the abundance of RE categories. We studied patterns of repeat dynamics within and across taxonomic groups by parsing RepeatMasker output tables and visualizing the distribution and abundance of major RE categories using custom python and R scripts.

*Comparison of assembly-based and cluster-based methods*
As an orthogonal approach to identifying repetitive elements with our assembly-based analysis, we explored genome repetitiveness and RE abundance with assembly-free approaches based on clustering of low-coverage short-read data as implemented in both dnaPipeTE v1.3.1 (Goubert et al. 2015) and RepeatExplorer2 (Novák et al. 2013). The former program relies on similar dependencies (e.g., RepeatMasker, Dfam, and Repbase) as our assembly-based approach and is thus well suited to exploring the effects of the clustering-based approach on RE estimates while reducing potential noise introduced by program-specific software and database dependencies. RepeatExplorer2 provides a comparison using a tool that uses a different underlying repeat database (i.e., Metazoa 3.0) and set of dependencies including TAREAN (Novák et al. 2017) which specializes in identification of satellite DNAs.

Samples for these analyses were chosen to both spread taxonomic representation across multiple insect orders and to minimize potential noise introduced by variation in technical details surrounding data generation. Briefly, we used NCBI metadata to identify samples for which the assembly was generated using long-read PacBio RSII chemistry (which was the category/sequencing chemistry of long-read assemblies that afforded the densest potential sampling), and for which metadata indicated that paired-end short-read Illumina data (specifically, either 100 or 150 paired-end data) were available for the same species as part of the same Bioproject. Of the 22 species that fit the above criteria we downloaded reads from NCBI's Sequence Read Archive and removed adapters/quality-trimmed raw reads using TrimGalore v0.6.3 (Krueger 2015). Trimmed, paired-end reads were mapped against indexed mitogenomes of the same species (or genus) using bwa mem 0.7.17 (Li 2013) (see additional methods for this step below). Reads which did not map to the mitogenome were identified with bam2fq.pl of FastQ Screen like tools (https://github.com/schellt/fqs-tools) and extracted with seqtk v1.3 command subseq (https://github.com/lh3/seqtk). Two samples for which we were unable to acquire a mitochondrial genome (either through de-novo assembly of the reads or finding the same species or a congeneric species on NCBI – see below) were excluded at this stage.

Paired, trimmed, mtDNA-filtered reads were then downsampled to 0.5x of the assembly length using seqtk and run using RepeatExplorer2 paired-end clustering with default settings and the taxon database Metazoa version 3.0. Single-end, trimmed, mtDNA-filtered reads were downsampled to 0.5x of the assembly length using seqtk and run in dnaPipeTE v1.3.1. For both analyses, runs that failed to complete were restarted using maximum allowed wall times which allowed all but five samples with failed runs to finish. We summarized overall repetitiveness and

RE abundance estimates from dnaPipeTE and RepeatExplorer2 as stacked bar charts and box plots ggplot2 (Wickham 2016) within R version 3.6.3 (R Core Team 2013).

*Filtering mtDNA from short read data*

Trimmed paired-end reads were mapped against the indexed mitogenomes of the same species (if a mitogenome of the same species was not available, we used a mitogenome of the same genus) using bwa mem 0.7.17 (Li, 2013) separately in unpaired mode. Alignments were printed to standard output in BAM format and sorted by leftmost coordinates using sort function with parameters -l 9 -O BAM -@10 of SAMtools v1.17 (Danecek et al., 2021). A list containing each read ID and the corresponding hit to the mitogenome was created using the script bam2fq.pl of FastQ Screen like tools for forward and reverse reads separately. The script classified unmapped reads as "No_hit". IDs of reads which did not map were extracted from these lists using grep -P "No_hit" | cut -f1. To restore paired information, it was checked, if both lists contain the ID and a final list of paired, unmapped reads was created using awk (cat ids_for_no_mt ids_rev_no_mt | sort | uniq -c | awk '$1==2{print $2}' > paired.ids_no_mt). Unmapped reads were extracted with the seqtk 1.3 command subseq. A list of mitochondrial genomes used for filtering can be found in on GitHub (https://github.com/johnssproul/Insect_REs) and Supplemental Materials.

*Correlation analyses*

We tested for correlations between RE abundance and a range of aspects for each genome assembly using R version 3.6.3 (R Core Team 2020). These included a comparison of total REs identified as well as specific classes (e.g., LINEs) versus the primary sequencing technology used (short- or long-reads). We also tested the relationship between the sequencing technology used and assembly length to assess the degree to which the technology might influence our findings. Next, we considered how genome assembly contiguity (as measured by contig N50) and completeness (as measured by BUSCO scores), compared to the same suite of RE total metrics (i.e., overall and classes). For all correlation analyses, we tested for normality in our data sets with a Shapiro-Wilk test and since the null hypothesis was rejected for all data sets ($P < 0.05$), we used Spearman's rank correlation tests.

*Repetitive elements and protein coding gene associations*

For all assemblies with ≥ 90% BUSCO gene content, we measured RE-gene associations (i.e., RE sequences inserted within or adjacent to protein-coding genes) following Heckenhauer et al (2021). Their study validated a new approach to quantifying RE sequences associated with BUSCOs. In some cases, the RE fragments are embedded within BUSCOs and in others REs with open reading frames that are immediately adjacent to BUSCOs are inadvertently classified by the BUSCO algorithm as being part of the BUSCO. They showed that quantifying such instances of RE sequences in BUSCOs can serve as a proxy for genome-wide RE-gene associations. We used a similar approach to Heckenhauer et al., except we adapted the protocol for higher throughput by using BLAST to identify BUSCOs containing RE sequences as opposed to using coverage read mapping coverage depth-based approach. Prior to moving forward with our modified approach, we performed a sanity check by directly comparing our BLAST-based methods to the coverage depth-based approach from Heckenhauer et al. (2022) in eight assemblies used in that study. We found estimates from our modified approach to be similar, but slightly more conservative (i.e., lower counts of RE-associated BUSCOs) in every comparison.

Our approach defined RE-associated BUSCOs as those which showed 10X the expected number of BLAST hits when used as a query against its genome assembly. To determine which

BUSCOs met this criterion, we stored each assembly as a custom BLAST database, used each BUSCO sequence as a query in a BLAST search against its respective assembly, and parsed the resulting output to make a table that summarized BLAST hit counts for each BUSCO query in a given assembly. We then sorted the BLAST output tables such that BUSCOs were ordered from lowest to highest BLAST hit counts and calculated the mean number of hits for the first three quantiles of the ordered data such that any BUSCOs with unexpectedly high blast hits (i.e., those residing in the fourth quartile of the data) would be excluded from the average calculation. We used this average as a baseline for expected BLAST hits, and any BUSCOs that had at least 10X the expected number of hits were considered “RE-associated BUSCOs”. Prior to settling on use of the first three quantiles as the fraction of data from which to calculate expected hit numbers, we plotted and visualized histograms of sorted BLAST hit counts for all assemblies and confirmed that BUSCOs with obvious hit inflation made up no more than 25% of all BUSCOs.

We searched for correlation between RE-associated BUSCOs and various measures in our data including sequencing technology (see above). We tested whether assembly artifacts might be driving the patterns of RE-associated BUSCOs by comparing detection trends in long-read vs short-read assemblies, with the expectation that if assembly artifacts were driving overall patterns we would see inflation of RE-associated BUSCOs in less contiguous short read assemblies.

*Investigating the effects of taxonomic sampling bias*

We investigated effects of taxonomic sampling bias on our understanding of REs in insects by analyzing the composition of the Repbase repository for RE sequences and the resulting impact on repeat annotation in our assemblies. We downloaded all Repbase database entries (01-27-2022 release) from https://www.girinst.org/repbase/. We used custom scripts to parse the insect database and quantify the taxonomic representation of insect orders and families included in our data set, as well as the rate of insect repeat submissions over time. We compared addition of new families to Repbase to data from Hotaling et al. (2021) which reported new family additions to Genbank.

In addition, we wanted to understand how the historical taxonomic bias in assemblies sequenced (in which some groups such as dipterans and hymenopterans are over-represented given their taxonomic diversity) would impact average assembly length and RE abundance values across the whole data set. To address this we randomly subsampled insect orders in proportion to their species abundance in order to re-frame general trends in assembly length and RE content without the bias introduced by over-represented lineages (e.g., Diptera, Hymenoptera, Lepidoptera). We used expected species abundance following Hotaling et. al (2021) and subsampled orders in proportion to ratios determined by expected values and calculated average assembly length and RE abundance based on the subsampled data.

### Supplemental Results

Given the uneven taxonomic sampling in our data set, we tested how normalizing the data set would impact overall trends we report by randomly sampling insect orders from our full data set in proportion to their estimated species richness (see Supplemental Materials). In essence, this corrected for the disproportionate abundance of species with small, relatively repeat-poor genomes which dominated early genomics projects. After reducing sampling bias in this way, we found that the average insect assembly length increased by 47.2% (from ~387 to ~569 Mb) and the average genomic proportion of LINEs, SINEs, DNA transposons, other repeats, and unclassified repeats showed all increased by 11.9–66.8%. This suggests that REs are more abundant in undersampled taxonomic groups.

### References


Danecek P, Bonfield JK, Liddle J, Marshall J, Ohan V, Pollard MO, Whitwham A, Keane T, McCarthy SA, Davies RM. 2021. Twelve years of SAMtools and BCFtools. *Gigascience* **10**: giab008.

Flynn, J.M., Hubley, R.*, Goubert, C., Rosen, J., Clark, A.G., Feschotte, C., Smit, A.F. (2020). RepeatModeler2: automated genomic discovery of transposable element gene families. *Proceedings of the National Academy of Sciences*, *117*(17), 9451-9457.

Goubert C, Modolo L, Vieira C, ValienteMoro C, Mavingui P, Boulesteix M. 2015. De Novo Assembly and Annotation of the Asian Tiger Mosquito (Aedes albopictus) Repeatome with dnaPipeTE from Raw Genomic Reads and Comparative Analysis with the Yellow Fever Mosquito (Aedes aegypti). *Genome Biol Evol* **7**: 1192–1205.

Heckenhauer, J., Frandsen, P. B., Sproul, J. S., Li, Z., Paule, J., Larracuente, A. M., ... & Pauls, S. U. (2022). Genome size evolution in the diverse insect order Trichoptera. GigaScience 11:giac011

Hoang, D. T., Chernomor, O., Von Haeseler, A., Minh, B. Q., & Vinh, L. S. (2018). UFBoot2: improving the ultrafast bootstrap approximation. *Molecular biology and evolution*, *35*(2), 518-522.

Hotaling, S., Kelley, J. L., & Frandsen, P. B. (2020). Aquatic insects are dramatically underrepresented in genomic research. *Insects*, *11*(9), 601.

Hotaling, S., Sproul, J. S., Heckenhauer, J., Powell, A., Larracuente, A. M., Pauls, S. U., ... & Frandsen, P. B. (2021b) Long-reads are revolutionizing 20 years of insect genome sequencing. *Genome biology and evolution*, evab138.

Katoh, K., & Standley, D. M. (2013). MAFFT multiple sequence alignment software version 7: improvements in performance and usability. *Molecular biology and evolution*, *30*(4), 772-780.

Kalyaanamoorthy, S., Minh, B. Q., Wong, T. K., Von Haeseler, A., & Jermiin, L. S. (2017). ModelFinder: fast model selection for accurate phylogenetic estimates. *Nature methods*, *14*(6), 587-589.

Kriventseva, E. V., Kuznetsov, D., Tegenfeldt, F., Manni, M., Dias, R., Simão, F. A., & Zdobnov, E. M. (2019). OrthoDB v10: sampling the diversity of animal, plant, fungal, protist, bacterial and viral genomes for evolutionary and functional annotations of orthologs. *Nucleic acids research*, *47*(D1), D807-D811.

Krueger F. 2015. Trim galore. *Wrapper Tool Cutadapt FastQC Consistently Apply Qual Adapt Trimming FastQ Files* **516**.

Kück, P., & Meusemann, K. (2010). FASconCAT: convenient handling of data matrices. *Molecular phylogenetics and evolution*, *56*(3), 1115-1118.

Kück, P., Meusemann, K., Dambach, J., Thormann, B., von Reumont, B. M., Wägele, J. W., & Misof, B. (2010). Parametric and non-parametric masking of randomness in sequence alignments can be improved and leads to better resolved trees. *Frontiers in zoology*, *7*(1), 1-12.

Li H. 2013. Aligning sequence reads, clone sequences and assembly contigs with BWA-MEM. *ArXiv Prepr ArXiv13033997*.

Minh, B. Q., Schmidt, H. A., Chernomor, O., Schrempf, D., Woodhams, M. D., Von Haeseler, A., & Lanfear, R. (2020). IQ-TREE 2: new models and efficient methods for phylogenetic inference in the genomic era. *Molecular biology and evolution*, *37*(5), 1530-1534.

Misof, B., & Misof, K. (2009). A Monte Carlo approach successfully identifies randomness in multiple sequence alignments: a more objective means of data exclusion. Systematic biology, 58(1), 21-34.

Novák P, Ávila Robledillo L, Koblížková A, Vrbová I, Neumann P, Macas J. 2017. TAREAN: a computational tool for identification and characterization of satellite DNA from unassembled short reads. *Nucleic Acids Res* **45**: e111–e111.

Novák P, Neumann P, Pech J, Steinhaisl J, Macas J. 2013. RepeatExplorer: a Galaxy-based web server for genome-wide characterization of eukaryotic repetitive elements from next-generation sequence reads. *Bioinformatics* **29**: 792–793.

Seppey, M., Manni, M., & Zdobnov, E. M. (2019). BUSCO: assessing genome assembly and annotation completeness. In *Gene prediction* (pp. 227-245). Humana, New York, NY.

Smit, AFA, Hubley, R & Green, P. *RepeatMasker Open-4.1.*2019 http://www.repeatmasker.org.

Swofford, D. (2003) PAUP, D. S. Phylogenetic analysis using parsimony (* and other methods), version 4. 2002. *Sinauer Sunderland, MA*.

Wickham H. 2016. *ggplot2: elegant graphics for data analysis*. Springer.

Zhang, C., Rabiee, M., Sayyari, E., & Mirarab, S. (2018). ASTRAL-III: polynomial time species tree reconstruction from partially resolved gene trees. *BMC bioinformatics*, *19*(6), 15-30.

### Supplemental Figures

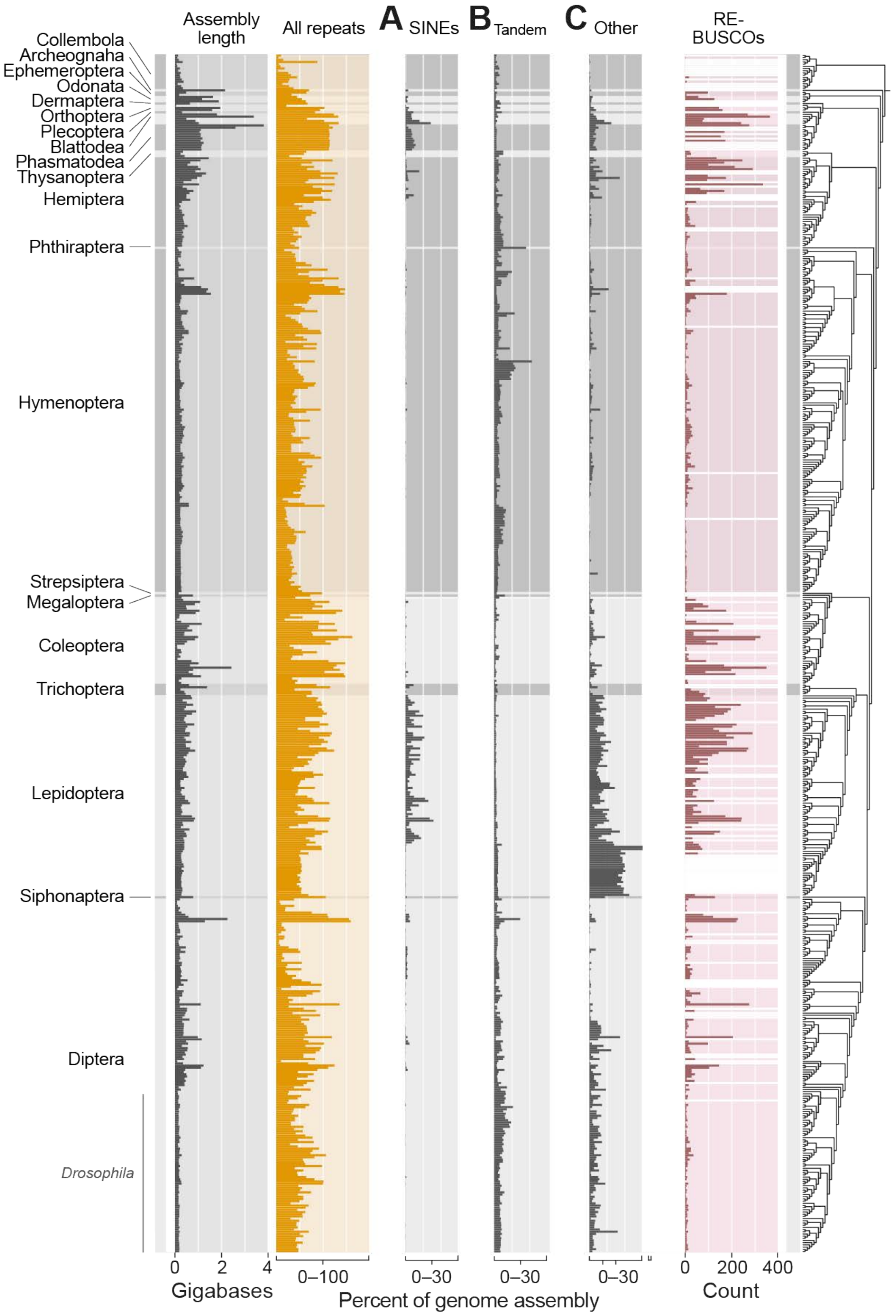


**Figure S1.** Genomic proportion of SINEs (A), tandem repeats (B), and other repeats (C) within the context of assembly length, total repeats, RE-associated BUSCOs, and phylogenetic relationships presented in Fig. 1.

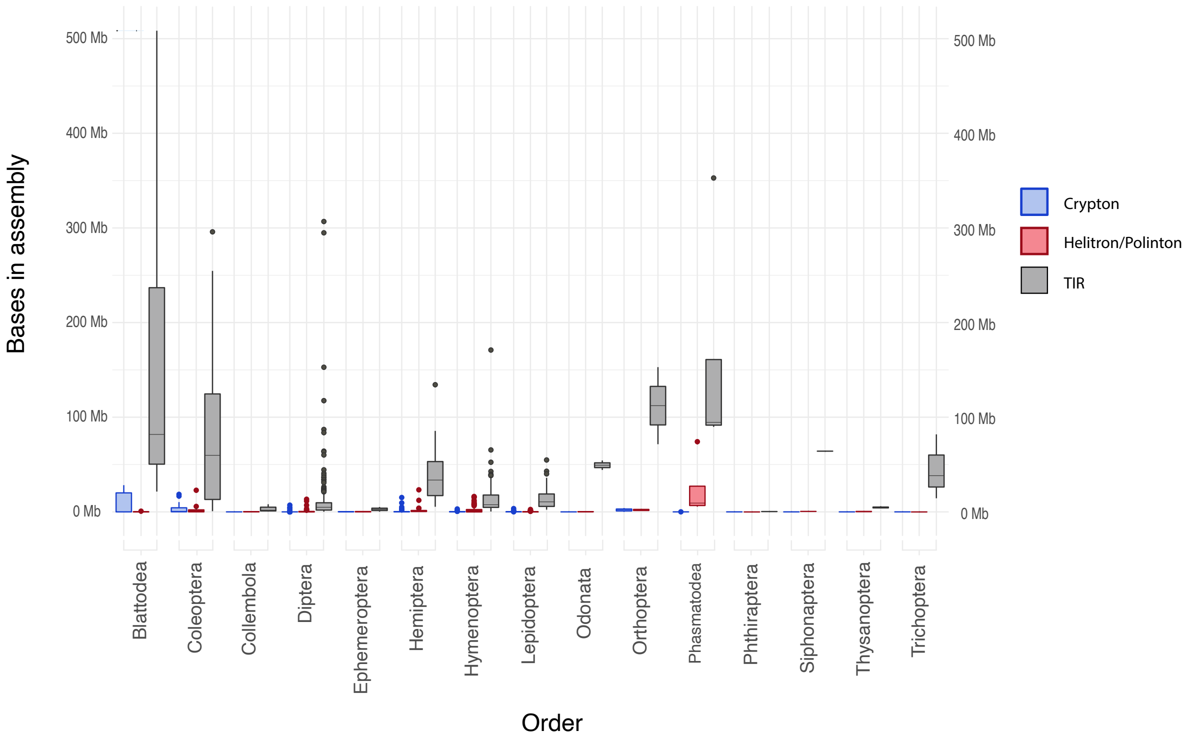


**Figure S2**. Abundance of major groups within Class II Transposable elements including Crypton, Helitron/Polinton, and TIR, summarized within each insect order that con-tained assemblies above the quality threshold of 90% BUSCO complete score.

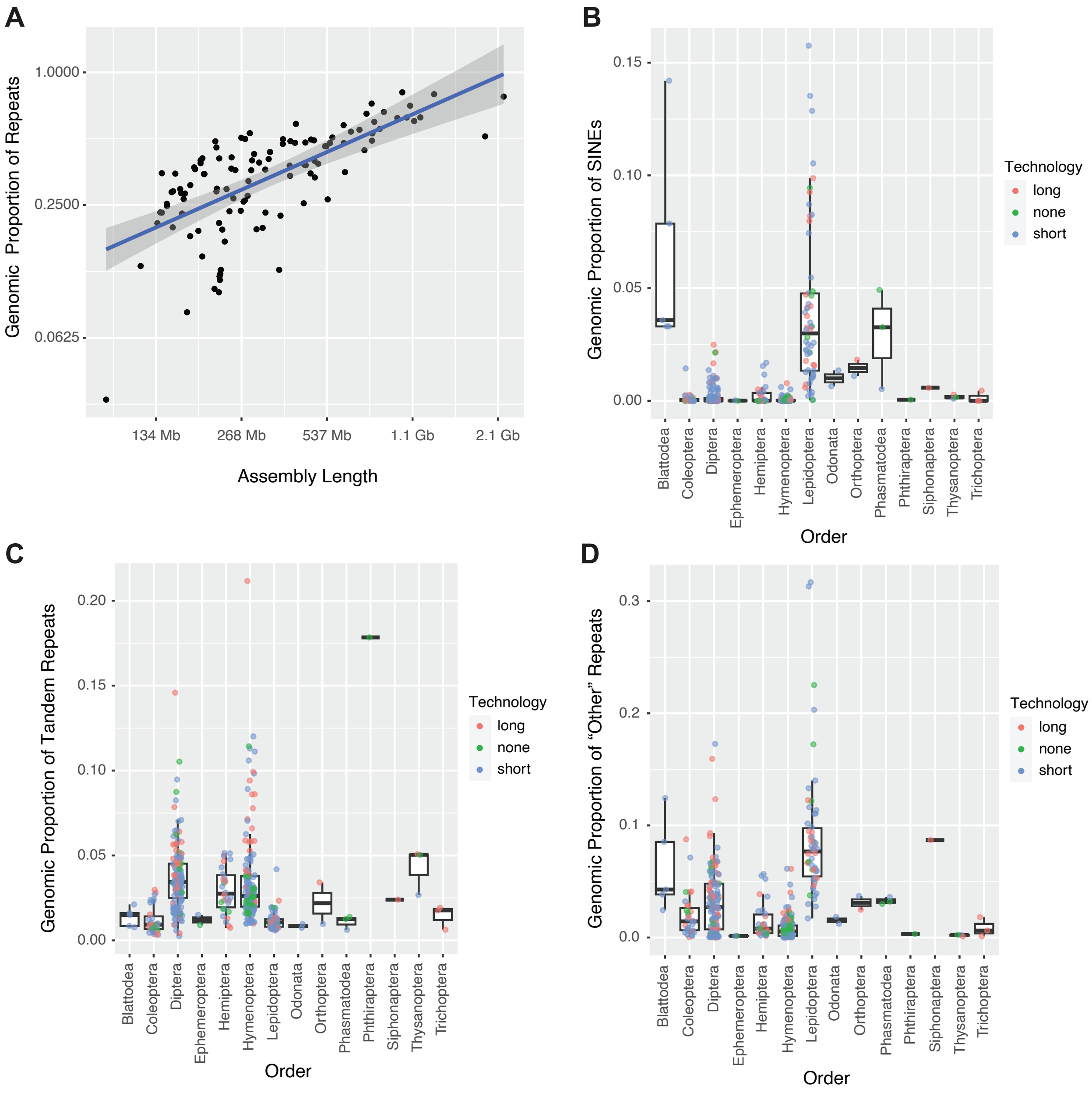


**Figure S3.** (A) Correlation between genomic proportion of repeats and assembly length to assess whether species with the largest genome assemblies show an inflection point at which genomic abundance of repeats falls below levels predicted by the trend line. The regression line is shown in blue and 99% confidence intervals are shown with gray shading. Analysis includes only long-read assemblies above the 90% complete BUSCO quality threshold. Genomic propotion of SINEs (B), tandem repeats (C), and "Other" repeats (D) with Helitrons accounting for the high abundance in some Lepidoptera species.

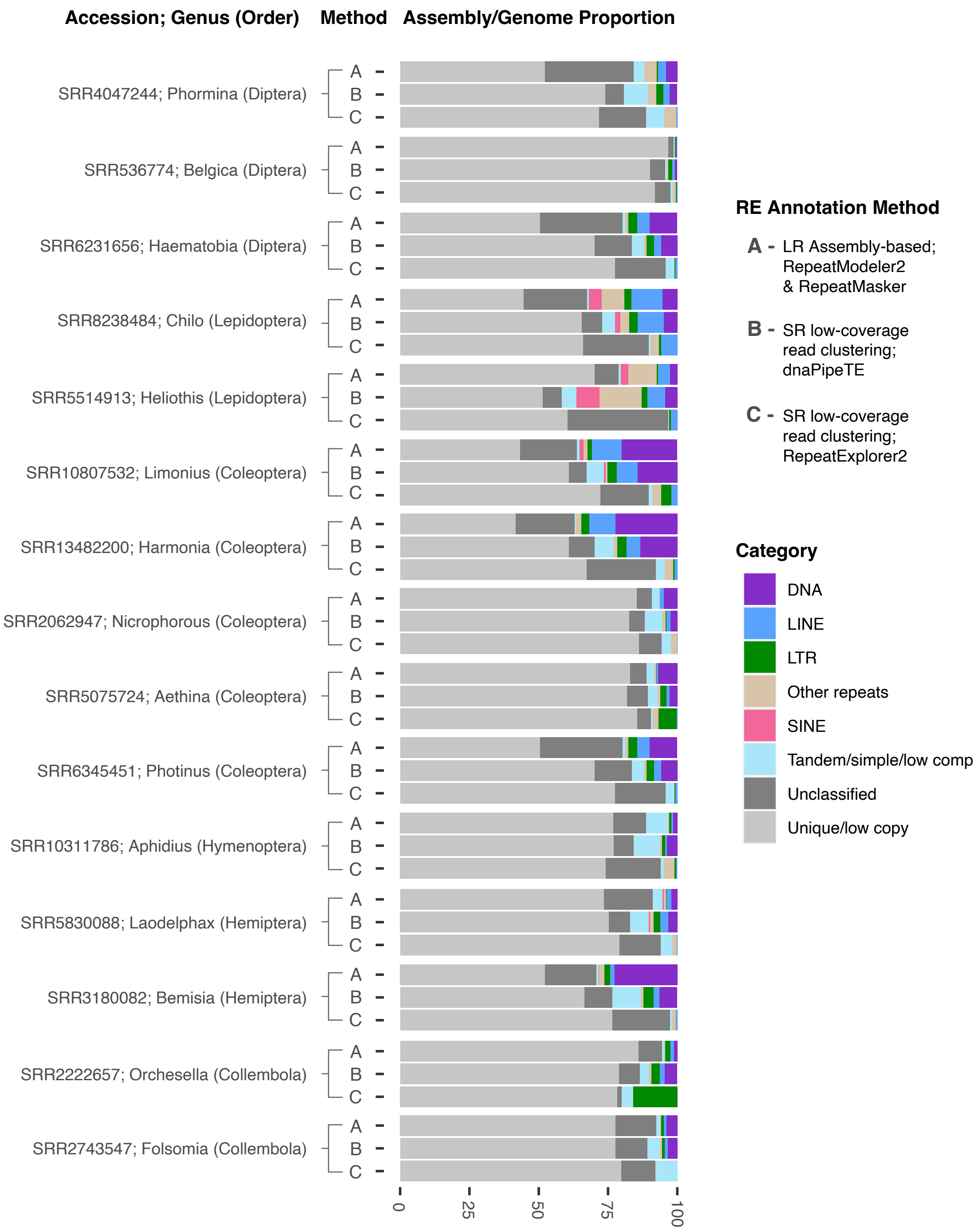


**Figure S4.** Comparison of assembly-based vs cluster-based RE detection in a subset of insect species. Method “A” used our standard assembly-based approach on long-read PacBio (RSII chemistry) assemblies using RepeatModeler2 and RepeatMasker for Re annotation. Methods “B” and “C” indicate dnaPipeTE and RepeatExplorer2 analyses, respectively. Both use assembly-free cluster-based analysis of short read Illumina (100 or 150 PE data) taken from the same studies for which PacBio assemblies were obtained. Note that “Other repeats” and “Tandem/simple/low comp categories are not entirely equivalent comparisons across analyses as each program differs slightly in RE categories reported in the output. For example, RepeatExplorer2 does not report simple or low complexity repeats and thus that category only represents tandem satDNA repeats for Method 3.

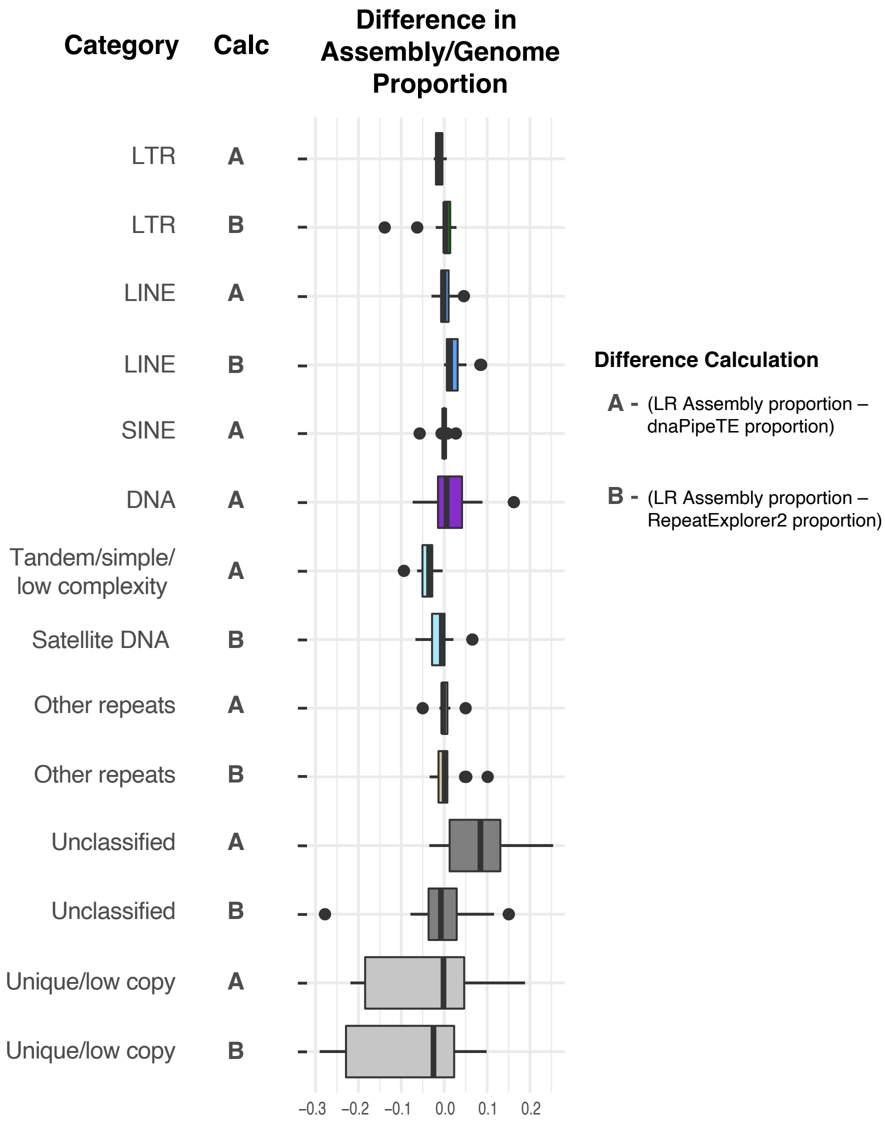


**Figure S5.** Difference in genome proportion in assembly-based vs cluster-based RE detection. For each repeat category the genomic proportion of REs detected in the long-read PacBio assemblies was subtracted from the proportion observed in the dnaPipeTE cluster-based approach (Calc "A"), and PacBio long-read assembly-based vs RE2 cluster-based approaches (Calc "B"). If box plots are distributed to the left of "0" on the x-axis, that repeat category was more abundant in the cluster-based analysis and less in the assembly-based. Right-distributed boxes followed the reverse trend. Note that not all comparisons could be made as RE2 lacks some of the categories of interest (hence, there is not a "B" for every instance of an "A").

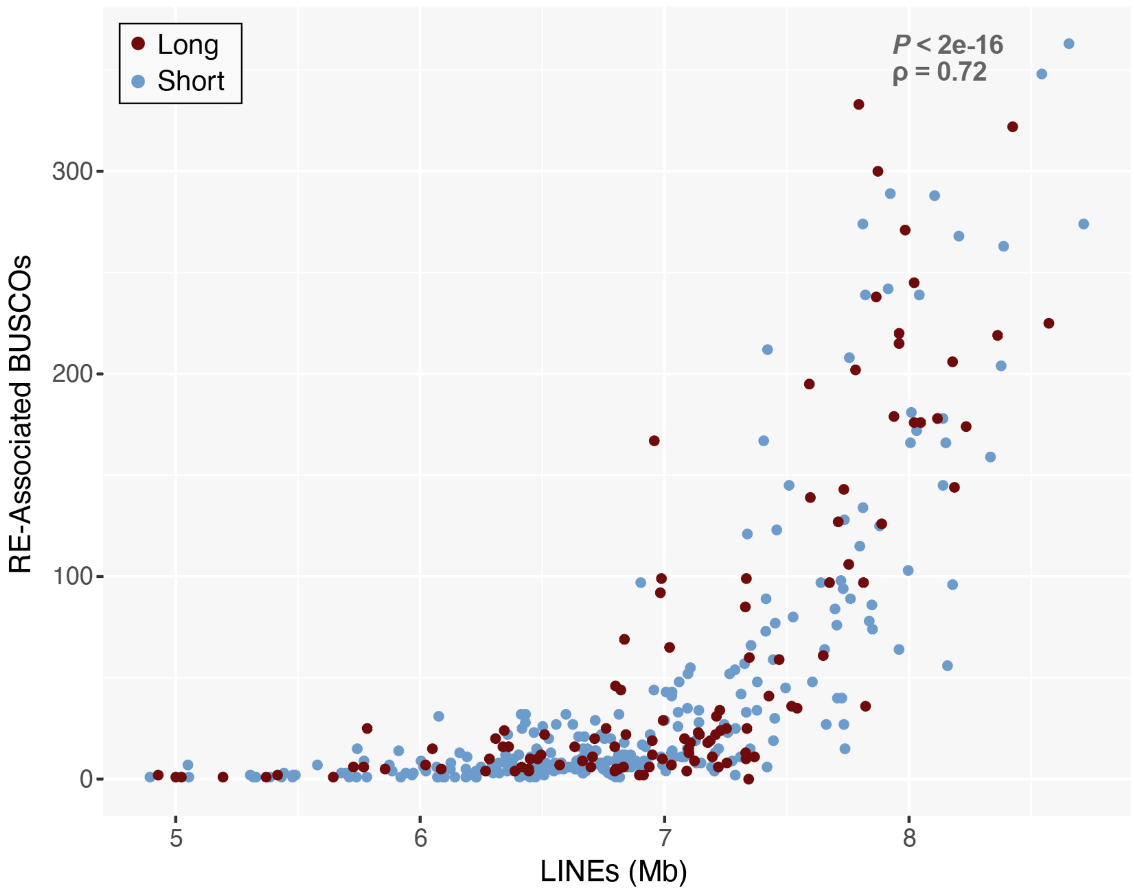


**Figure S6.** Spearman correlation for the number of RE-associated BUSCOs and bases of LINEs across insect genomes, color-coded by sequencing technology.

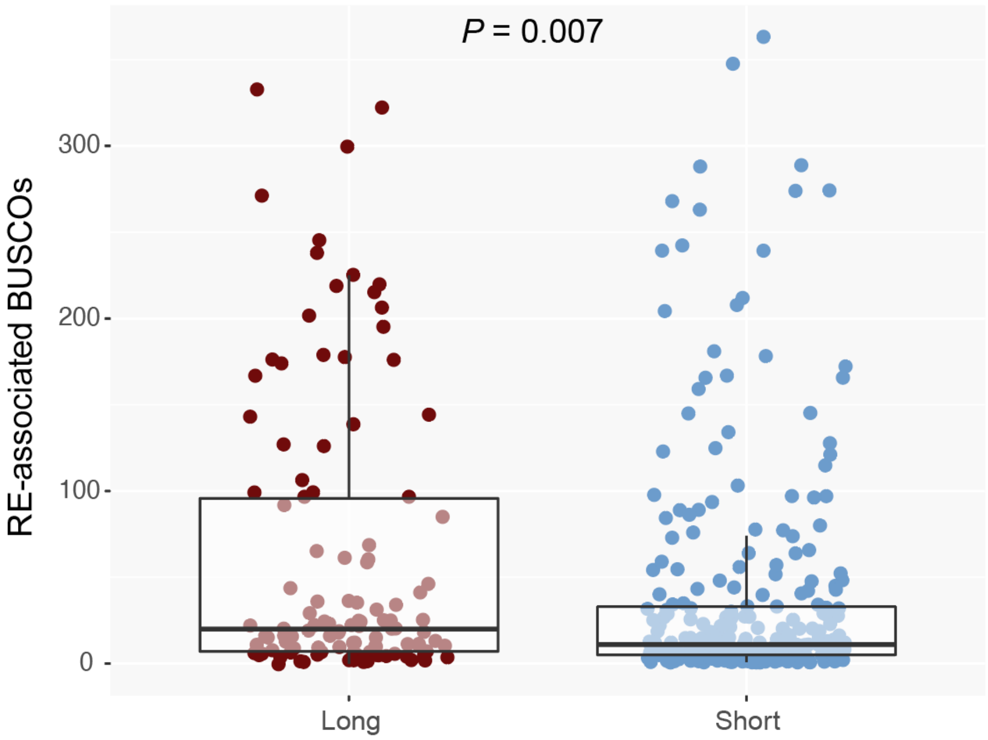


**Figure S7.** A comparison of the number of RE-associated BUSCOs identified versus the sequencing technology used. Significance was assessed with Welch two-sample T-tests.

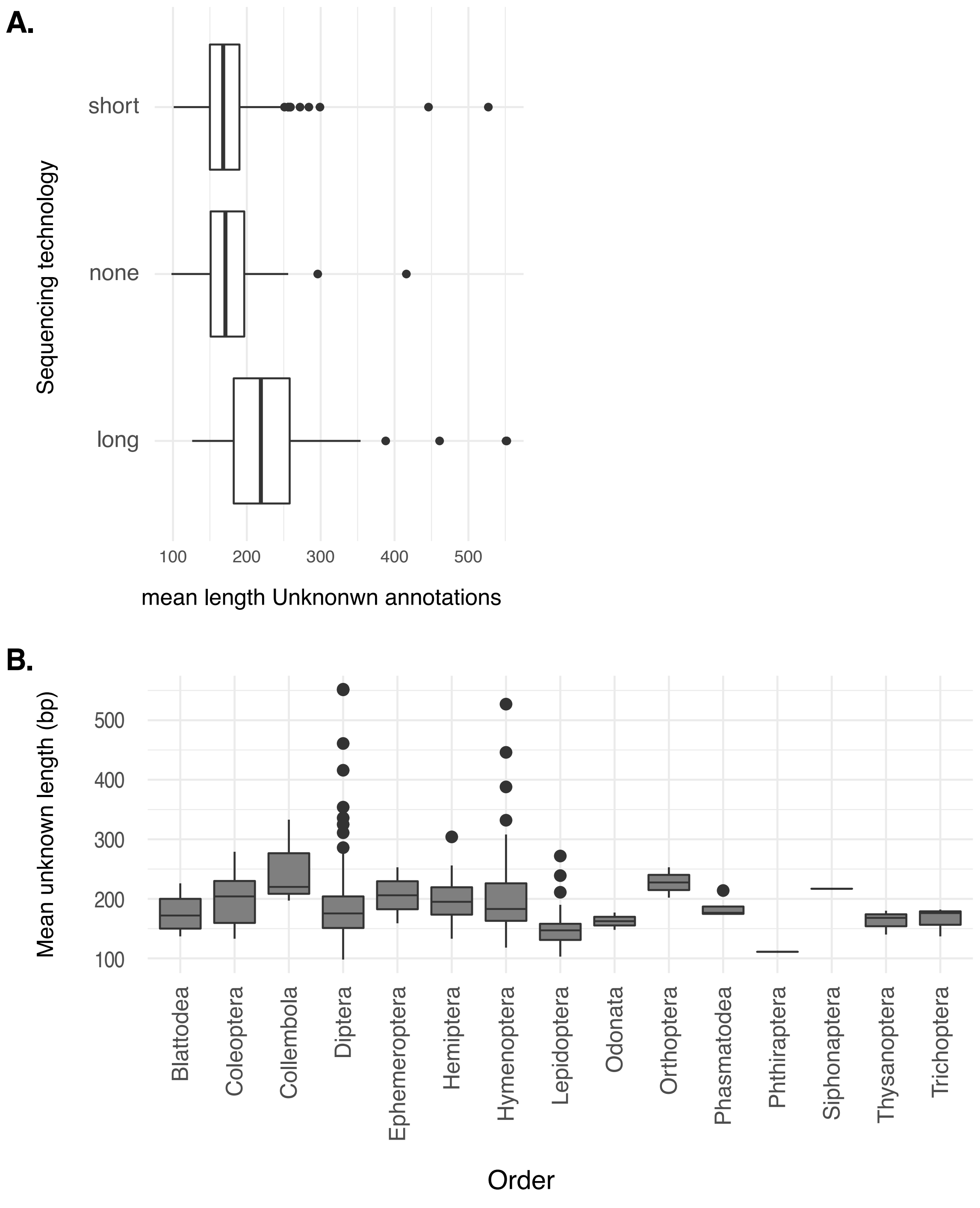


**Supplemental Figure S8**. Length distribution of unknown repeats based on sequencing technol-ogy (A), and broken down by insect order (B).
